# Sustained immune activation suppresses planarian regeneration

**DOI:** 10.64898/2026.05.17.725704

**Authors:** Noam Hendin, Omri Wurtzel

## Abstract

Tissue injury immediately triggers immune defenses to prevent infection, a process that can paradoxically interfere with repair. Yet, how some organisms resolve this tension to fully regenerate remains poorly understood. Planarians, flatworms capable of regenerating any body part, offer a unique model for studying how robust immunity coexists with extensive regenerative capacity. Here, we show that the planarian immediate injury response is dominated by the robust upregulation of immune and stress-related genes, demonstrating that defense mechanisms are intrinsically wired into wound sensing. By uncoupling immune activation from tissue injury using exposure to heat-inactivated bacteria, we found that immune stimulation alone induced a transcriptional program mirroring central aspects of the early injury response. Prolonged immune activation led to progressive, host-driven tissue lysis that was fully reversible upon removal of the stimulus. Single-cell profiling identified distinct epidermal and phagocytic subpopulations as the central mediators of this “defense-first” response. Furthermore, we identified *foxF-1*-regulated phagocytes as critical drivers of immune resolution, as suppressing *foxF-1* markedly increased vulnerability to noninfectious immune challenge. Finally, we demonstrated that sustained immune hyperactivation delayed regenerative progression by approximately 50%. Together, our findings establish the resolution of immune activity as a critical prerequisite for regeneration and define sustained immune activation as a fundamental constraint on tissue repair.

## Introduction

Tissue injury triggers a cascade of physiological processes essential for survival ^1–4^. Following tissue damage, wounds must be rapidly sealed, pathogens eliminated, and debris cleared, while maintaining an environment permissive to tissue repair ^3,5,6^. Across metazoans, this response triggers a broadly conserved transcriptional program organized into temporally distinct phases, beginning with inflammation ^2,7–9^. Wounding rapidly induces immediate-early genes, such as *fos-1, jun-1* and *early-growth response genes (EGRs)* ^9–12^, followed by activation of inflammatory cytokines and stress-response factors that recruit immune cells and prime local defense ^13–16^. The cytotoxic environment created by inflammatory signaling is crucial for destroying pathogens, but simultaneously damages the host tissues ^17–20^, conflicting with the requirements for tissue repair ^21^. In non-regenerating species, injury often provokes a hyperactive, persistent inflammation that typically results in fibrosis or chronic wounds, or severe pathologies like sepsis ^17,21–23^. Conversely, in regenerating organisms, new tissue growth is associated with more moderate and transient activation of inflammatory immune signaling ^24–28^.

The inverse relationship between robust immunity and regeneration has been described in different animals. For example, in *Xenopus*, the maturation of the adaptive immune system during metamorphosis coincides with the loss of the ability to regenerate limbs ^23,29–31^. Similarly, neonatal mammals, which have an under-developed immune system incapable of mounting a strong inflammatory response, are able to regenerate skin and cardiac tissues, an ability that is lost during postnatal maturation ^32–34^. These observations imply an evolutionary trade-off between regenerative capacity and an advanced immune response ^14,24,33^.

The freshwater planarian *Schmidtea mediterranea* is a powerful model for dissecting how immune programs operate in a regeneration-compatible manner. Planarians are capable of regenerating any body part, yet they are highly resistant to a broad range of pathogens and environmental stressors, demonstrating an effective immune defense that does not compromise regenerative repair ^35–38^. Furthermore, because planarians lack an adaptive immune system, they allow for the precise isolation of innate immune dynamics during regeneration.

Planarian injury activates a generic transcriptional wound response ^9,39,40^, characterized by the rapid, transient induction of stress genes and various immune-associated gene homologs, such as *TRAFs, C-type lectins, PGRPs, PIM1* and *PIM2* ^41–46^. While the transcriptional profile of this response has been described ^9,39,41^, the extent to which physical injury activates a bona fide innate immune program – and whether this program suppresses or promotes regeneration initiation – remains unresolved.

In this work, we characterize the planarian inflammatory and immune responses, and demonstrate the dramatic consequence of prolonged inflammation on regeneration. We first show that an innate immune-like transcriptional program is intrinsically wired into the immediate injury response, driven primarily by epidermal cells and phagocytes. To isolate this immune response from the confounding effects of physical wounding, we challenged animals with heat-killed bacteria. We found that immune activation alone induced a state of suppressed regeneration, where the persistent inflammatory signaling delayed both general tissue regrowth and the production of eye cells and sensory neurons by approximately 50%. Moreover, we identified *foxF-1*^+^ phagocytes as potential regulators of this response, demonstrating that their suppression renders animals hypersensitive to immune challenges. Together, our findings establish that the resolution of immune activity is a fundamental prerequisite for planarian regeneration, revealing that a sustained immune response actively suppresses inherent regenerative potential.

## Results

### Injury triggers an immune program that precedes new tissue formation

To define early transcriptional responses to injury, we analyzed a published RNA-seq time course of planarian injury ^41^, focusing on the initial 24 hours post-amputation (hpa). Differential expression analysis detected 349 upregulated genes following injury, which resolved into six distinct temporal modules based on their expression patterns (Fig. 1A-B; Table S1; FDR < 0.05; Fold-change ≥ 2; Methods). Temporal gene expression patterns correlated to functional programs. Early clusters (peaking at 3–6 hpa) showed rapid, transient induction of immediate-early transcription factors (e.g., *fos-1*, *jun-1*), stress-response genes (e.g., *HSP*s, *STIP1*), and immune-related homologs (e.g., *TRAFs*, *PIM1*, *DUSP10*) (Fig. 1A-B; Table S1)^46–48^. Conversely, late-induced clusters (16–24 hpa) included regeneration-associated genes, including tissue polarity, differentiation, and patterning factors, such as *wntP-3*, *sp9*, and *ATOH1* (Fig. 1A; Table S1) ^49–51^. Intermediate clusters, induced by 6 hpa and sustained through 24 hpa, contained immune-associated genes encoding lectin-like factors, extracellular matrix remodelers (e.g., *ADAMTS1*), and regulators of DNA repair and oxidative stress ^52,53^ (Fig. 1A-B; Table S1). Because animals were maintained in a mostly pathogen-free environment, these results indicated that tissue damage alone is sufficient to activate an innate immune-like transcriptional program.

**Figure 1.**
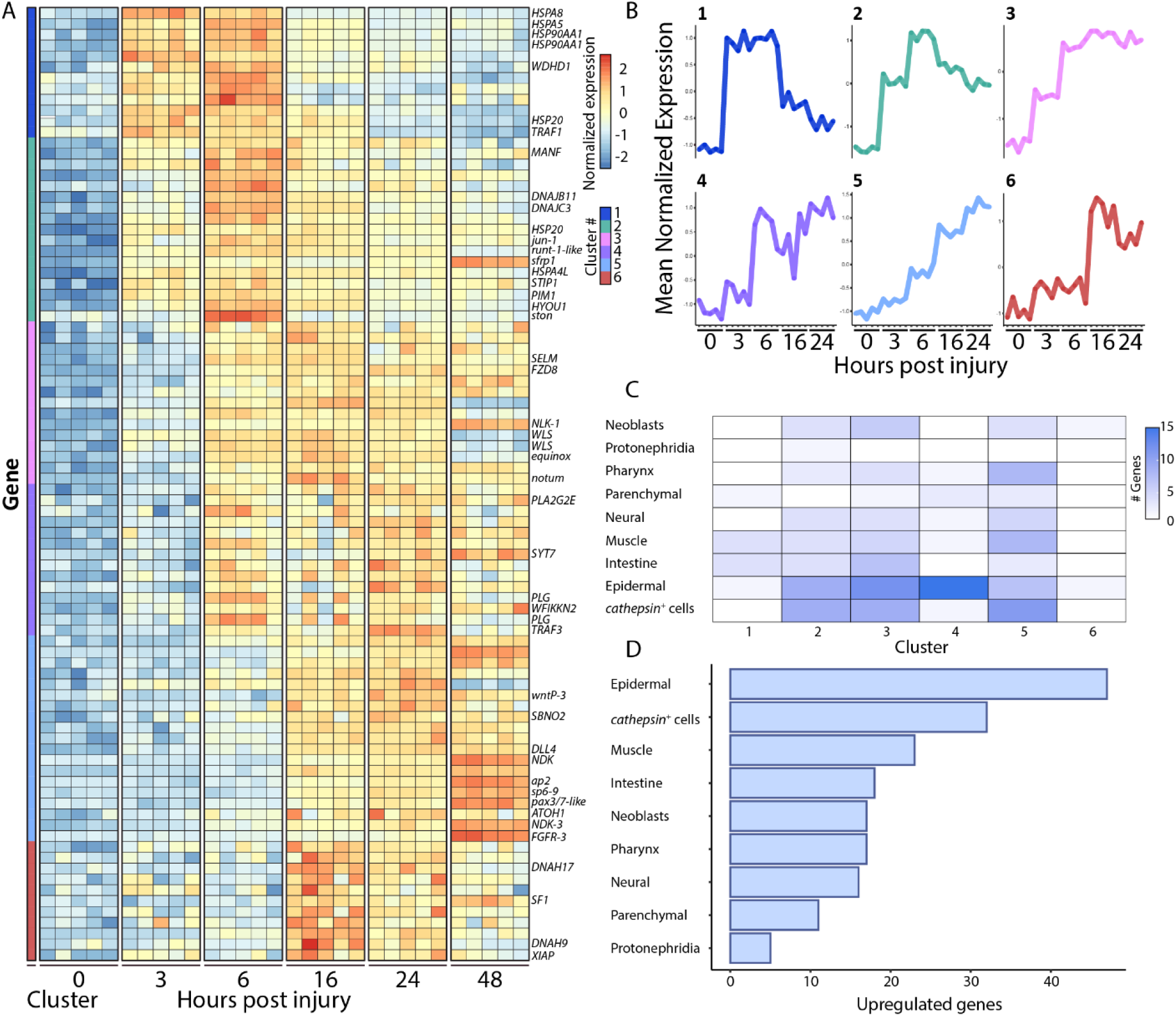
Transcriptional changes following amputation and cell types involved in the planarian immediate injury response. (A) Heatmap showing upregulated gene expression following injury of representative genes of each cluster (blue and red, low and high gene expression, respectively; expression is shown as row normalized z-scores). Clustering analysis determined six groups of differentially expressed genes based on their expression patterns over time (Table S1; Methods). (B) Line graphs plot the mean normalized expression of the genes within each of the six temporal gene expression clusters across the sampled time points. (C) Heatmap showing the number of upregulated genes from each cluster in specific planarian cell types ^57^. Color intensity represents the number of genes enriched in each cell lineage. (D) Bar chart quantifies the total number of upregulated wound-response genes assigned to each cell type, demonstrating that epidermal and *cathepsin*^+^ cells are major drivers of the immediate transcriptional response.

Of the injury-induced genes, we identified 66 immune-related genes, distributed across all six clusters (Table S1; Methods). These genes were classified as immune activators, suppressors, or both, based on injury context and supporting literature (Table S1; Methods). Overall, immune activators and suppressors were induced in similar numbers (29 and 24 genes, respectively) across the response, with comparable proportions across clusters 1-5 (Table S1). Examination of this distribution suggested some cluster-specific differences in immune-associated gene composition. Cluster 3 contained the highest number of immune genes (19) and the greatest relative representation of suppressor-associated factors (∼47%). By contrast, clusters 1, 4, and 5 showed a higher relative representation of activator-associated genes, with cluster 5 exhibiting the strongest bias (∼64% of immune genes classified as immune activators). Notably, by 48 hpa, during which new tissue is generated ^54,55^, this immune-like transcriptional program was largely downregulated (Table S1), marking a transition from an initial inflammatory-like state to the onset of tissue regeneration.

### Epidermal and phagocytic cells drive the planarian immediate wound response

We next analyzed the cell types mediating the immediate injury response. While prior studies established epidermal and muscle lineages as the major hubs of wound-induced positional information signals ^9,39,41^, our analysis identified planarian phagocytes (*cathepsin*^+^ cells) and epidermal cells as the primary drivers of the early immune-like response. These two populations expressed the greatest number of wound-induced genes (Fig. 1C-D). Lineages responsible for positional information and tissue growth, such as muscle cells and neoblasts (planarian pluripotent stem cells) ^9,56^, exhibited critical but fewer transcriptional changes during this immediate window (Fig. 1C-D; Table S1).

Mapping specific gene clusters to cell types revealed distinct temporal contributions. The earliest peaking cluster (1) was enriched for stress-response genes (e.g., *HSP*s, *HYOU1*), which are broadly expressed and not associated with specific tissue types (Fig. 1A-C) ^57^. Mid-early peaking clusters (2–3), however, contained immune activating factors (*TRAFs*, *jun-1*, and *fos-1*), and were associated primarily with *cathepsin*^+^ and epidermal cells (Fig. 1A-C; Table S1). Furthermore, cluster 4 genes were almost exclusively expressed by epidermal cells; a module containing canonical immune activating genes but lacking patterning factors (Fig. 1C; Table S1). Clusters 3 and 5 displayed similar temporal expression trends, but also contained patterning-associated genes. While gene expression in these clusters was driven by multiple cell types, epidermal and phagocytic cells were dominant (Fig. 1C). Muscle and pharyngeal cells notably increased their expression of cluster 5 genes, aligning with the high concentration of patterning genes in this module (Fig. 1A-C; Table S1). Together, these data indicated that epidermal and phagocytic lineages successively activated transcriptional programs at the early phases of the wound response.

### Non-infective immune activation recapitulates central features of the early wound response

The immune system shapes tissue repair outcomes ^24,58^. Our analysis of the planarian injury response indicated that an immune response was activated prior to new tissue growth. To determine the role of the planarian immune system in the tissue repair process, we uncoupled immune activation from physical tissue damage. We achieved this by exposing intact planarians to heat-killed *Escherichia coli*, acting as a non-pathogenic immune stimulant (Fig. 2A; Methods). Because inactivated bacteria present antigens but cannot actively infect tissues and cause disease ^59–62^, this approach allowed the isolation of immune responses from disease- and wounding-related transcriptional changes.

**Figure 2.**
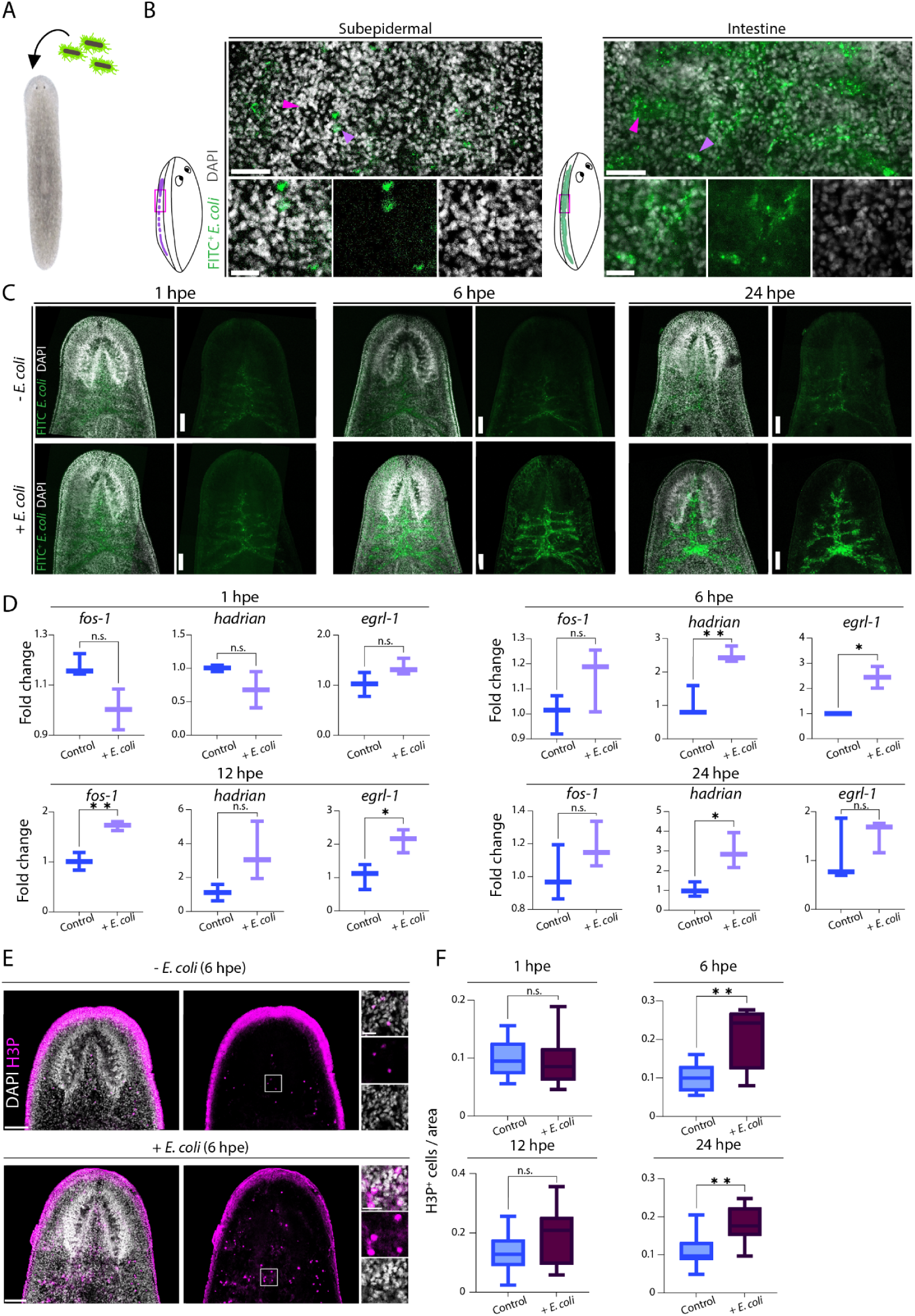
Exposure to heat-killed *E. coli* partially recapitulates injury response in planarians. (A) Experiment outline; FITC-labeled heat-killed *E. coli* (green) was added to planarian water for exposure. (B) FITC-labeled heat-killed *E. coli* (green) was detected in the sub-epidermal parenchyma (left) and intestine (right); arrows (magenta) indicate major structures (left - ventral nerve cord; right - intestinal lumen); purple arrows indicate intracellular bacteria. Scale: 50 and 25 µm, top and bottom panels, respectively. Diagrams indicate tissue regions shown (ventral nerve cords, left; intestine, right), labeled in purple and green, respectively. (C) Detection of FITC-labeled heat-killed *E. coli* (green) in internal planarian tissues. Top panels: without *E. coli* (unexposed), bottom panels: with *E. coli* (exposed); for each time point, right panels display FITC-channel only. Bacteria were not detected at 1 hpe. Scale: 100 µm. (D) qPCR quantification of injury-induced gene expression following exposure to heat-killed *E. coli*; three biological replicates were used for each condition (Methods); Y-axis indicate fold change compared to control. P-values were calculated using unpaired two-tailed Student’s t-test; error bars indicate minimal and maximal values. (E) Representative images of H3P labeling of worms unexposed (top) and exposed (bottom) to bacteria at 6 hpe. Scale: 100 µm; H3P^+^ cell in magenta; white squares indicate region of high-magnification shown in right panels. Scale: 25 µm. (F) H3P^+^ cell counts normalized to area (Methods) following exposure to heat-killed *E. coli*. Y-axes indicate normalized counts of H3P^+^ cells; horizontal line indicates median; error bars indicate minimal and maximal values. Number of worms counted per group: 1 hpe control group: n = 5; 1 hpe + *E. coli*: n = 10; 6 hpe control: n = 9; 6 hpe + *E. coli*: n = 9; 12 hpe control: n = 10; 12 hpe + *E. coli*: n = 10; 24 hpe control: n = 10; 24 hpe + *E. coli:* n = 9. P-values were calculated by unpaired two-tailed t-test (** p <0.01, * p <0.05), horizontal line indicates median.

We first confirmed that planarians internalized the inactivated bacteria. Following exposure to FITC-labeled heat-killed *E*. *coli*, confocal imaging detected no internalization at 1 hour post-exposure (hpe) (Fig. 2B-C; Methods). However, by 6 and 24 hpe, FITC signal localized predominantly within the intestinal lumen, as well as in the subepidermal parenchyma, and was detected intracellularly (Fig. 2B-C). This broad distribution confirmed that the inactivated pathogen reached internal tissues where it could potentially trigger a systemic host response.

To test if a host response was activated by the heat-killed bacteria, we analyzed the expression of canonical stress-activated genes *fos-1* and *egrl-1,* and the planarian-specific wound-induced gene *hadrian* ^9,39,40^. Worms were isolated at 1, 6, 12 and 24 hours following exposure to heat-killed *E. coli*, and RT-qPCR was used to quantify gene expression at each time point (Methods). Consistent with the timing of bacterial internalization, this qPCR analysis showed no significant expression changes at 1 hpe, but revealed significant upregulation of the three genes at 6, 12, and 24 hpe (Fig. 2D). This indicated that these injury-induced genes also function directly in the planarian innate immune program.

We next assessed whether immune stimulation induces stem cell proliferation, a second hallmark of the planarian injury response ^54^. Physical wounding typically triggers two distinct waves of neoblast division: an initial body-wide generic response peaking around 6 hours post-injury (hpi), and a subsequent localized wave starting around 48 hpi specific to missing tissue ^54^. Following exposure to heat-killed *E. coli*, phospho-histone H3 (H3P) labeling revealed a significant increase in systemic stem cell division at 6 and 24 hpe, but not at 1 or 12 hpe (Fig. 2E-F; Fig. S1A-C; Methods). Importantly, worms did not display visible damage or compromised viability at this point. This wave-like proliferative response has been reported following infection with live *Candida albicans* ^63^, further suggesting it is part of the planarian defense program. Ultimately, these data showed that immune activation using non-infective particles was sufficient to induce the fundamental transcriptional and cellular hallmarks of the early wound response, even in the complete absence of physical injury, further showing a wiring of the two responses.

### Sustained immune activation leads to tissue damage and systemic lysis

Following injury, the planarian immune and stress transcriptional program resolves prior to 48 hpa, immediately preceding the onset of regenerative patterning^39^ (Fig. 1A; Table S1). Because continuous inflammatory signaling disrupts stem cell dynamics and drives severe pathologies across animals ^17,26,64,65^, this rapid downregulation suggests that resolving the inflammatory phase functions as a biological prerequisite for active tissue repair. To test if unresolved immune activation directly interferes with planarian tissue maintenance, we exposed animals to continuous immune stimulation, beyond 24-hours, to prevent the resolution of the immune response.

Strikingly, extending the exposure to the heat-killed bacteria was detrimental and induced progressive tissue damage, starting with dorsal lesions, progressing to head regression and ultimately resulting in complete lysis (Fig. 3A). Lethality typically occurred approximately three days post-exposure (in worms sized ∼5-6 mm), with smaller animals (∼2 mm) succumbing earlier, approximately 48 hours following exposure (Fig. 3A). Removing the immune stimulant halted further tissue damage and permitted full recovery, which corresponded with a decline in host-response gene expression (Fig. S2A-B). Because the bacteria were non-viable, this damage was likely the result of a cytotoxic, host-driven hyper-inflammatory response rather than active infection. This chronic response was also accompanied by elevated stem cell mitotic activity at 48 hours, indicating active neoblast participation in prolonged host defense (Fig. 3B-C). Furthermore, exposing planarians to various other live and heat-killed bacteria (including *P. aeruginosa* and *B. subtilis*) consistently triggered similar tissue lysis, demonstrating a generic, sepsis-like host response to sustained bacterial sensing (Fig. S2C).

**Figure 3.**
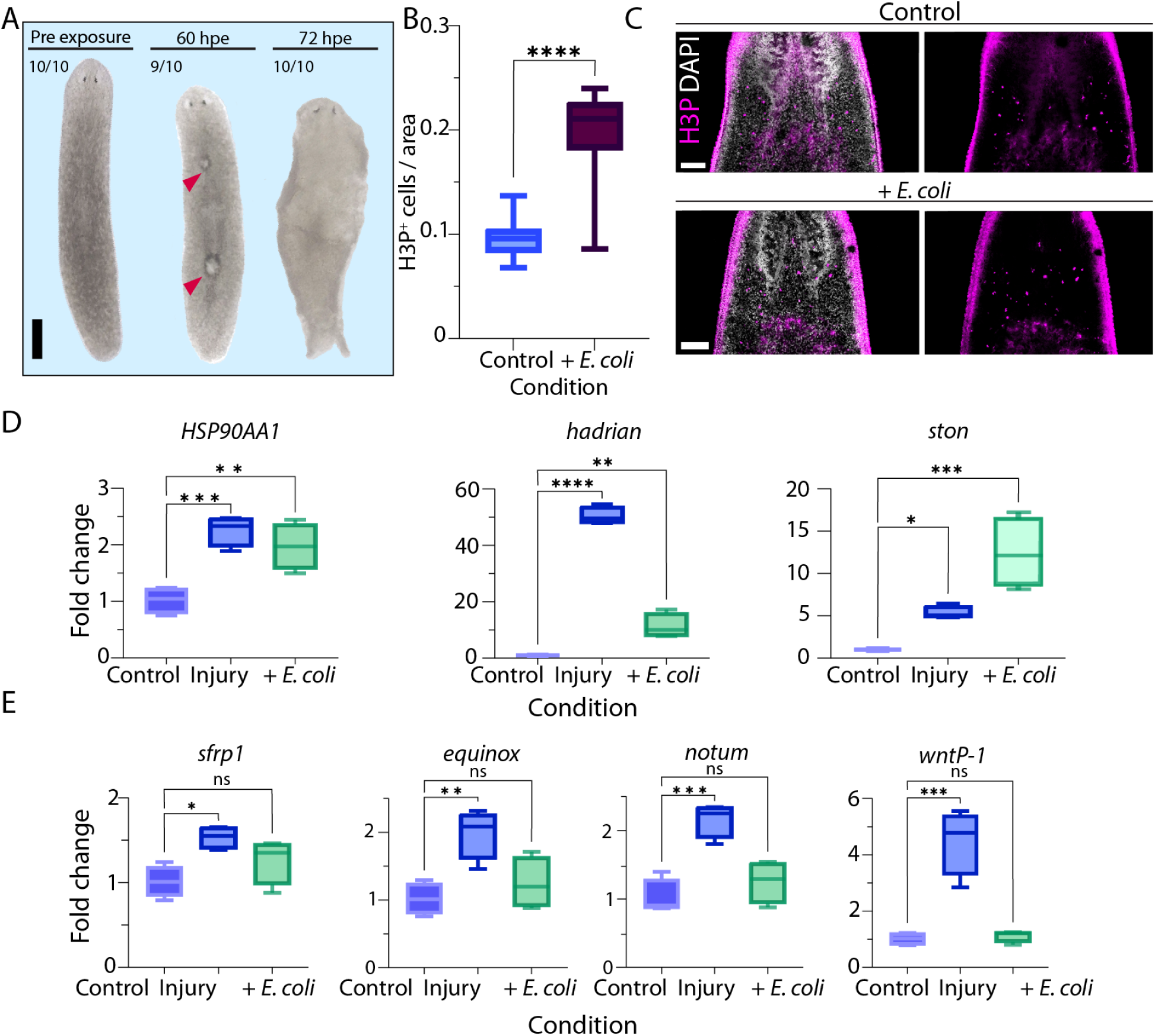
Heat-killed bacteria activate a subset of injury-induced genes and induce worm lysis. (A) Prolonged exposure to heat-killed *E. coli* causes progressive tissue damage. By ∼60 hpe, 90% of the worms developed dorsal lesions (red arrowheads); by ∼3 dpe all worms underwent lysis. (B) H3P^+^ cell counts of unexposed worms (control) vs. worms exposed to heat-killed *E. coli* at 48 hpe; p-values were calculated by unpaired two-tailed Student’s t-test (**** p <0.0001), horizontal line indicates median; error bars indicate minimal and maximal values. (C) Representative images of H3P^+^ nuclei in control (top) and treated (bottom) worms. Scale: 100 µm. (D-E) RT-qPCR quantification of injury-induced genes not associated with patterning (D) and patterning-related injury-induced factors (E). Y-axes indicate fold change, horizontal lines indicate median. P-values were calculated by one-way ANOVA, followed by Holm-Šídák’s multiple comparisons test (**** p <0.0001, *** p <0.001, ** p <0.01, * p <0.05), horizontal lines indicate median; error bars indicate minimum and maximum values.

To determine if this bacterial exposure specifically induced the immune module of the wound response without triggering the regenerative module, we compared the expression of classical injury-induced genes following either physical wounding or heat-killed *E. coli* exposure. By 12 hours, both physical injury and bacterial exposure significantly upregulated the stress-related gene *HSP90AA1*, as well as the epidermal genes *hadrian* and *ston* (Fig. 3D). By contrast, while physical injury strongly induced patterning and differentiation genes (*sfrp-1*, *equinox*, *notum*, and *wntP-1*), bacterial exposure did not activate these regenerative factors (Fig. 3E). This transcriptional divergence demonstrated that exposure to inactivated bacteria selectively triggered the innate defense program while excluding tissue-patterning pathways. Therefore, the induction of early wound-induced genes like *hadrian* and *ston* by immune stimulation indicated they function specifically within the planarian defense response.

### An innate immune-like transcriptional program is activated in the absence of physical injury

Prolonged exposure to heat-killed pathogens induced inflammatory lesions and lysis in the absence of physical injury (Fig. 3A). To understand the transcriptional basis of this sepsis-like outcome, we profiled gene expression in animals exposed to heat-killed *E. coli* at 20 and 40 hpe (Fig. 4A; Methods). This profiling identified a distinct immune and stress module containing canonical immune-associated genes (e.g., *PIMs*, *TRAFs*, C-type lectins, *HSP*s, and *EGRs*) without concurrent activation of patterning or differentiation genes (Fig. 4A; Table S2) ^11,46,47,66–68^. Comparison with live *E. coli* exposure at 40 hpe revealed high transcriptional similarity (Fig. S3A-B, Table S3; R = 0.86), confirming that inactivated bacteria elicit a bona fide pathogen-defense program.

**Figure 4.**
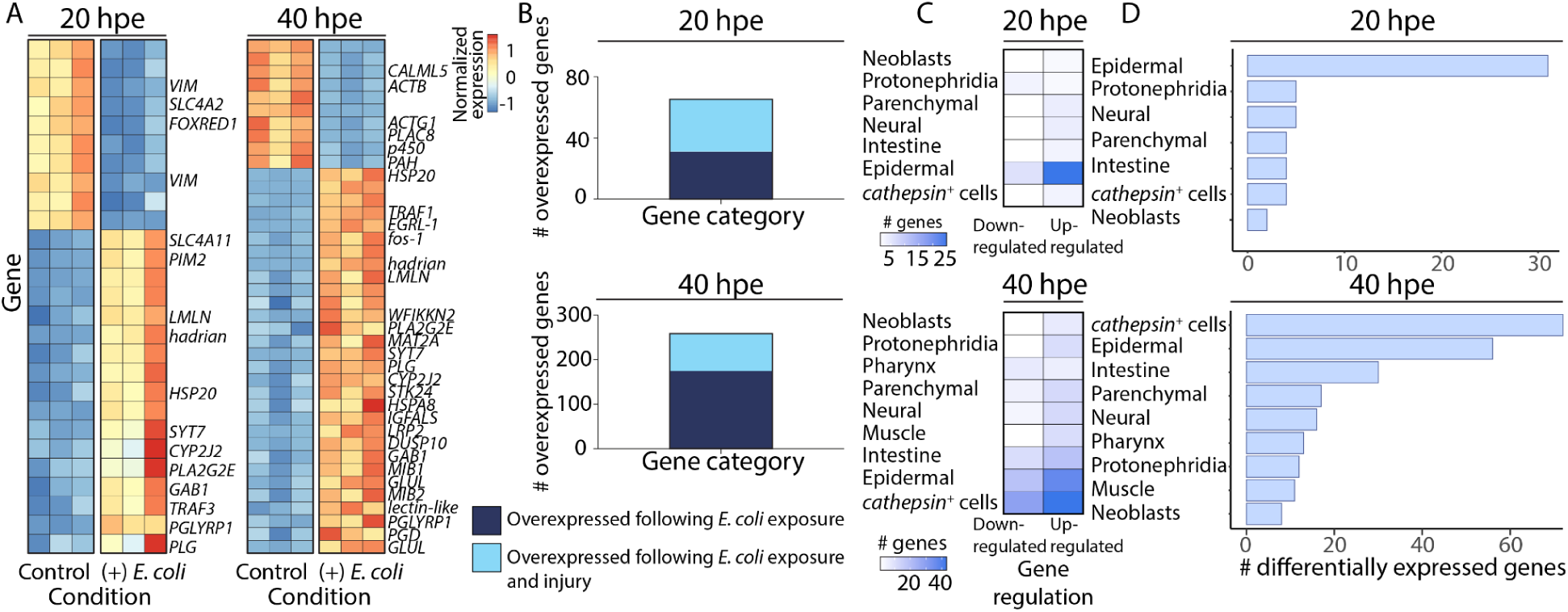
Exposure to heat-killed *E. coli* activates a bi-phasic innate immunity-like program in planarians. (A) Heatmaps displaying differentially expressed genes at 20 (left) and 40 (right) hours post-exposure (hpe), representing the early and late stages of the response, respectively (blue and red, low and high gene expression, respectively; expression is shown as row normalized z-scores). (B) Bar plots comparing injury and host response gene expression; dark blue bars showing the number of genes uniquely induced by the host response, and light blue bars display up-regulated genes in both the host and injury responses; top: number of genes upregulated at 20 hpe compared to injury time course showing 34/65 genes are common to both responses (Fisher’s exact test p = 6.71x10^-33^); bottom: number of genes upregulated at 40 hpe compared to injury time course showing 83/258 genes are common to both responses (Fisher’s exact test p = 3.37x10^-59^). (C) Heatmaps show the number of differentially expressed genes across cell types at 20 hpe (top) and 40 hpe (bottom); color intensity indicates the number of genes enriched in each cell lineage. (D) Bar graphs quantifying the total number of DEGs of the host response assigned to each cell type, at 20 hpe (top) and 40 hpe (bottom).

At 20 hpe, the transcriptional response was restricted to 91 differentially expressed genes (DEGs; log fold-change ≥ |0.9|; FDR < 0.05; Table S2). This gene set represented an initial response, characterized by the presence of genes encoding immune activators, such as *phospholipase A2* (*PLA2G2E*) and *synaptotagmin VII* (*SYT7*). By 40 hpe, this response extended to 410 DEGs (Table S2). To characterize the functional shift over time, we annotated transcripts as immune activators or immune suppressors (Methods). Immune-related transcripts constituted 30.8% (n = 12/39) and 30% (n = 67/223) of all annotated DEGs at 20 and 40 hpe, respectively (Table S2). Notably, the fractions of immune activators and suppressors were comparable between time points (activators: 50% and 52.2% at 20 and 40 hpe, respectively; suppressors: 33.3% and 37.3% at 20 and 40 hpe, respectively; Table S2). Genes that were classified as having both immunosuppressive and immune activating functions (n = 7) were not included in this analysis.

Comparing this host response to the physical injury dataset revealed significant overlap in upregulated genes (Fig. 4B, Table S2; Methods). At 20 hpe, 52.3% of the upregulated genes (34/65) were shared with the injury response (Fig. 4B; Table S2; Fisher’s exact test p = 6.71x10^-33^). This intersection indicated that the early response to immune stimuli comprises an intrinsic module of the generic injury response program ^39^. By 40 hpe, this overlap decreased to 32.2% (Fig. 4B; Table S2; 83/258 genes; p = 3.37x10^-59^). This decreased overlap indicated that prolonged exposure triggered a distinct late-phase immune program, separate from the generic wound response.

Mapping the DEGs to the planarian cell-type transcriptome atlas identified the cellular basis for this shift (Fig. 4C-D; Methods) ^57^. At 20 hpe, epidermal cells were the primary responders in the early-stage host response, expressing the majority of DEGs (Fig. 4C-D). This occurred despite bacterial particles showing predominant localization in the intestine at this stage (Figs. 2B-C). By 40 hpe, however, phagocytes (*cathepsin*^+^ cells) emerged as the central mediators, accompanied by sustained epidermal expression (Fig. 4C-D). This cellular shift demonstrated a dynamic spatial response to continuous immune stimulation. The delayed activation of phagocytes suggested they either propagate the cytotoxic host response or function to promote its resolution ^27,69^.

### Single-cell RNA sequencing reveals that specific epidermal and phagocytic cell populations mediate the host response

To determine which cell populations drive the immune response, we performed single-cell RNA sequencing 40 hours after exposure to heat-killed bacteria (Methods). We analyzed a total of 17,649 cells (8,901 treated, 8,748 control), resolving 34 clusters that encompassed all major planarian cell types and states (Fig. 5A; Fig. S4A; Table S4-S5; Methods). The relative abundance of these clusters remained comparable between conditions (Fig. S4B), indicating that the immune response relied on transcriptional shifts within existing populations rather than the expansion or depletion of specific cell types. By comparing gene expression between control and heat-killed bacteria-exposed samples, we assigned transcriptional responses to specific cell types (Fig 5B-C; Table S5). We identified both cell type-specific responses and genes differentially expressed across multiple lineages (Fig 5B-C). This high-resolution analysis enabled us to identify epidermal and *cathepsin*^+^ cell populations as central mediators of this stage of the host response (Fig 5D).

**Figure 5.**
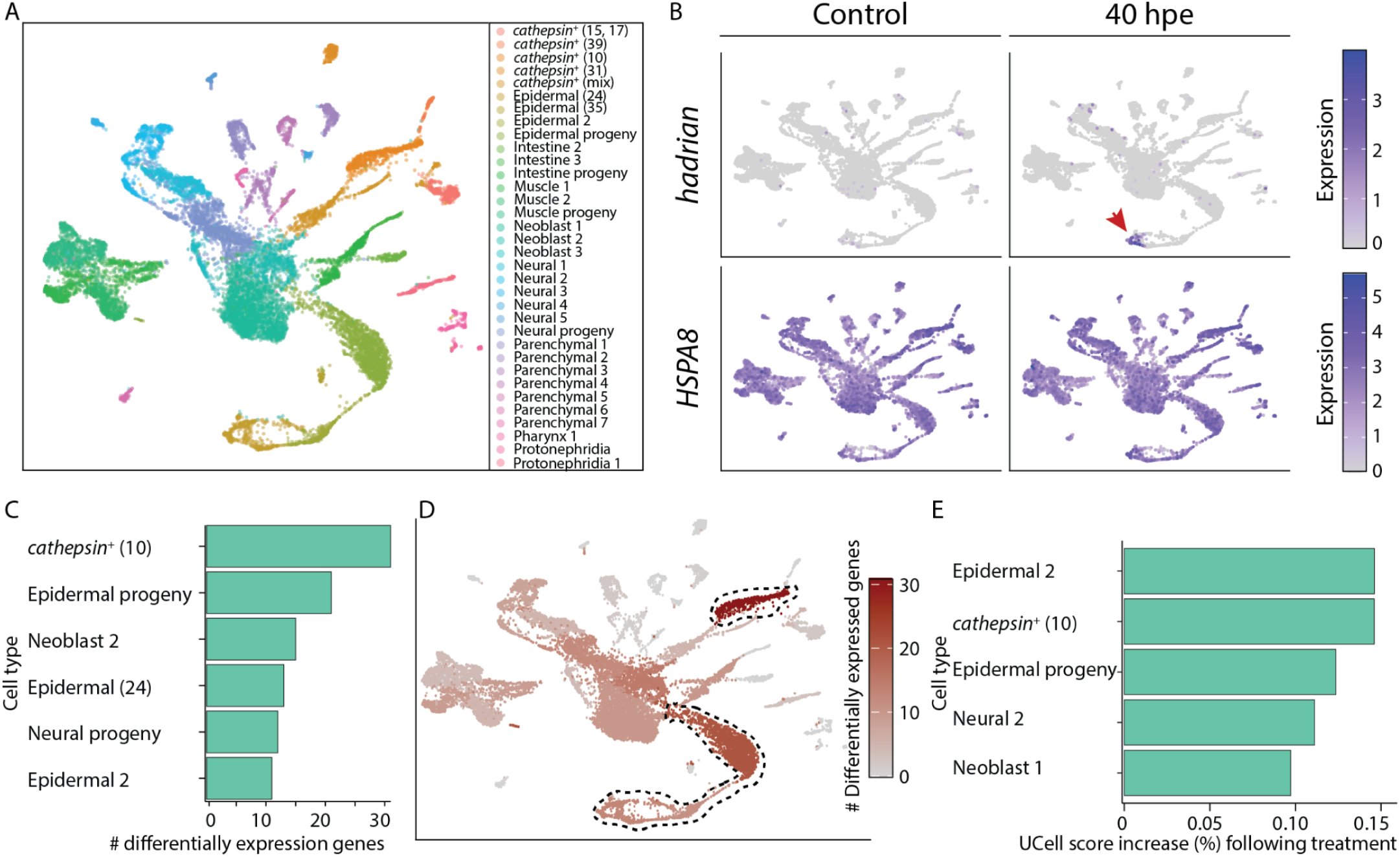
scRNA-seq analysis reveals sub-populations of *cathepsin*^+^ and epidermal cells as primary responders to bacterial exposure. (A) UMAP projection of sequenced cells, annotated by distinct cell types and lineages based on the planarian cell atlas ^57^. (B) Feature plots comparing the spatial expression patterns of *hadrian* and *HSPA8* between control samples and samples at 40 hours post-exposure (hpe) to bacteria. The red arrowhead indicates the localized expression of *hadrian* within a specific cell cluster (epidermal, 24) following exposure. By contrast, the overexpression of *HSPA8* was broad and spanned multiple cell types. (C) Bar plot quantifying the number of differentially expressed genes across the top responding cell clusters. A *cathepsin*^+^ cluster exhibited the greatest number of transcriptional changes following exposure to heat-inactivated bacteria. (D) UMAP projection with cells colored by the number of differentially expressed genes per cluster. The dashed lines highlight the subpopulations of *cathepsin*^+^ and epidermal cells showing the greatest number of overexpressed genes. (E) Mean UCell scores for immune-related genes between heat-killed bacteria treated and control conditions were used to calculate the relative increase in UCell score. Shown is UCell score increase for cell populations having a significant increase (FDR < 0.01; Methods).

To quantify these shifts, we measured expression enrichment scores (UCell) ^70,71^ of the immune-related gene set, detected in the bulk RNA-seq, across all clusters (Table S2; Methods). We observed a robust, cluster-specific increase in immune-related gene transcriptional programs, with the largest mean increases occurring in epidermal cluster 2 (∼14.7%) and *cathepsin*^+^ cluster 10 (∼14.5%) populations (Fig. 5E; Figure S4A). A minor increase, in Neoblast 1, was detectable, but more modest (∼9.7%), suggesting that while stem cells sense the immune challenge, the primary biological response is concentrated in these two differentiated lineages. These results identified specific epidermal and *cathepsin*⁺ cell populations as the central responders of this stage of the host response.

Interestingly, a specific *cathepsin*^+^ subpopulation exhibited the most extensive transcriptional changes (Fig 5C-D, S4A; Table S5), acting as the primary responding cell type at this time point. This cell population distinctly expresses *granulin* ^57^ (Fig. S4C), a gene encoding a conserved immune modulator that has both pro- and anti-inflammatory functions ^72,73^. Altogether, these data established that the planarian immune response relied on transcriptional reprogramming, mediated largely by epidermal and *granulin*-expressing *cathepsin*^+^ cell populations.

### *foxF-1* suppression accelerates immune-driven tissue lysis

By 40 hpe, *cathepsin*⁺ phagocytes emerged as the dominant transcriptional responders (Fig. 4C-D, Fig. 5C-D), prompting us to investigate whether they promote the cytotoxic response or drive its resolution. Our RNA-seq data revealed that during this late stage of the host response, these cells expressed similar numbers of immune activators and suppressors (Table S2; Table S4). To functionally distinguish between these two roles, we targeted the *cathepsin*⁺ lineage using RNA interference (RNAi) against *foxF-1*, a broad master regulator of *cathepsin*⁺ subtypes ^57,74^.

As *foxF-1* is a regulator of visceral muscle (in addition to *cathepsin*^+^ phagocytes) ^74^, we applied a mild suppression to target the phagocyte lineage without causing muscle deterioration (Methods). Following *foxF-1* RNAi, planarians exhibited the characteristic depigmentation phenotype, consistent with effective target silencing (Fig. 6A) ^74,75^. *foxF-1* knockdown was validated by RT–qPCR (Fig. 6B), and retention of visceral musculature integrity was confirmed by immunolabeling for a *foxF-1*-regulated muscle marker (Fig. 6C; Methods). Notably, labeling of phagocytic cells using the broad *cathepsin* marker *CSTL2*⁺ revealed no significant reduction in this population following this modest *foxF-1* suppression (Fig. 6D-E).

**Figure 6.**
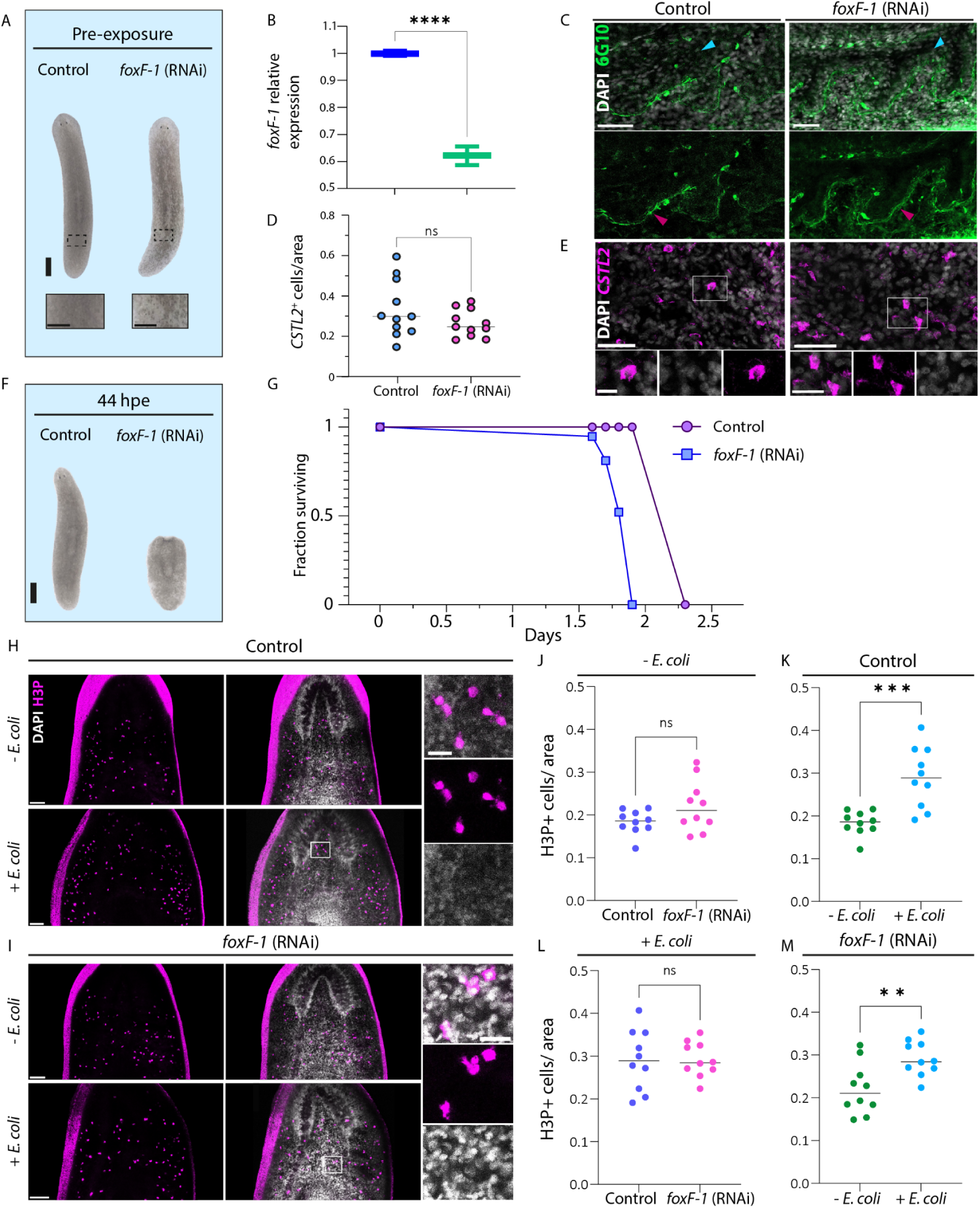
*foxF-1* suppression accelerates deterioration under sustained, non-pathogenic immune activation. (A) Representative images of control and *foxF-1* (RNAi) animals 9 days post-feeding (pre-exposure to immune stimulant). *foxF-1* (RNAi) animals exhibited characteristic depigmentation, consistent with previous reports ^74,75^. Scale bars: 0.5 mm (main) and 0.25 mm (magnified squares). (B) RT-qPCR quantification of *foxF-1* gene expression in control compared to *foxF-1* (RNAi) animals at 7 days post-feeding (Methods). (C) Immunofluorescence of *foxF-1* lineage-derived internal muscle (green) at 9 days post-feeding. Muscle surrounding the intestine (red arrows) showed comparable intensity between control and *foxF-1* (RNAi) animals, indicating preserved muscle integrity. Blue arrows indicate intestinal lumen. Bottom panels display isolated muscle labeling. Scale bar: 50 µm. (D-E) Quantification (D) and representative FISH images (E) of *CSTL2*^+^ cells at 9 days post-feeding, showing no significant difference in cell count between conditions. Y-axis shows *CSTL2*^+^ cells per area (Methods). Bottom panels show high-magnification images. Scale bars: 50 µm (top) and 25 µm (bottom; magnified). (F) Comparison of animals at 44 hpe to heat-killed *E. coli*. *foxF-1* (RNAi) animals showed head-regression and lethality (n = 10/14) compared to controls, which remained intact and viable (n = 12/12). Scale bar: 0.5 mm. (G) Kaplan-Meier survival curve of *foxF-1* (RNAi) vs. control (RNAi) worms following immune challenge. *foxF-1* suppression (blue squares) resulted in rapid mortality starting at 38 hpe (n=3/14), reaching 0% survival by 46 hpe, compared to 55 hpe in control animals (purple circles). (H-I) Representative images of H3P⁺ cells (magenta) in control (RNAi) (H) and *foxF-1* (RNAi) (I) animals, without (top) and with (bottom) heat-killed *E. coli* present. Scale: 100 µm; right panels show a magnification of H3P^+^ nuclei. Scale: 25 µm. (J-M) Normalized H3P^+^ cell counts (mitoses per area) in *foxF-1* (RNAi) and control worms, before and after heat-killed *E. coli* challenge. P-values were calculated using unpaired two-tailed Student’s t-test. (J) No significant difference between homeostatic control and *foxF-1* (RNAi) worms (n=10 each group). (K, M) Both control and *foxF-1* (RNAi) animals significantly increased mitoses after 20h heat-killed *E. coli* exposure, although the response was slightly attenuated in *foxF-1* (RNAi). (L) Comparison at 20h post-exposure showed similar total mitotic counts between conditions (n=10 each). P-values were calculated by unpaired two-tailed Student’s t-test; * p <0.05, ** p < 0.01, *** p <0.001, **** p < 0.0001; error bars indicate minimal and maximal values.

We then exposed the *foxF-1* (RNAi) animals to heat-killed *E. coli*. The *foxF-1* (RNAi) animals exhibited accelerated tissue damage, undergoing head regression and complete lysis approximately 24 hours earlier than control RNAi animals (Fig. 6F-G). Unstimulated *foxF-1* (RNAi) planarians remained viable for weeks with visibly intact musculature (Fig. S5), confirming that this accelerated mortality resulted directly from a compromised response to the immune challenge rather than poor physical conditions. Together, these data demonstrated that the *foxF-1*-regulated phagocyte program acts to attenuate the inflammatory host response, though this buffering capacity ultimately failed under continuous immune stimulation.

### Immune-induced stem cell proliferation is unaffected by *foxF-1* suppression

We next quantified the neoblast response following suppression of phagocyte gene expression, as our analysis identified phagocytes as major contributors to the host response (Fig. 4C-D, Fig. 5C-D). Importantly, in planarians, *foxF-1*-lineage phagocytes associate closely with neoblasts and function as a stem cell niche ^74,76^. We tested whether *foxF-1*-specified phagocytes regulated the proliferative neoblast response induced by exposure to heat-killed bacteria (Fig. 2E-F; Fig. 3B-C). We hypothesized that decreased phagocyte gene expression via *foxF-1* suppression would in turn decrease neoblast divisions during immune stimulation. We administered *foxF-1* or control dsRNA to planarians (Methods). Ten days after the final feeding, we separated the animals into unexposed and exposed groups (exposed or unexposed to heat-killed *E. coli*; Fig. 6H-M). We then fixed the planarians 20 hours post-exposure and labeled them for phospho-histone H3 (H3P) to quantify mitotic activity (Methods) ^54^.

Unexposed planarians showed no significant difference in H3P⁺ cell counts between control and *foxF-1* RNAi conditions (Fig. 6J). Exposure to heat-killed *E. coli* significantly increased mitotic activity in both control and *foxF-1* RNAi animals relative to their respective unexposed baselines (Fig. 6K, M). Direct comparison of exposed *foxF-1* and control RNAi planarians, however, showed no significant difference in H3P^+^ cell labeling (Fig. 6L). These results demonstrated that *foxF-1* suppression did not alter homeostatic or immune-induced stem cell proliferation. Consequently suggesting that *foxF-1*-specified phagocytes do not regulate the stem cell proliferation associated with immune activation. Instead, these data implied that this phagocyte subpopulation specifically attenuated tissue lysis during the cytotoxic host response, whereas the observed increase in stem cell proliferation represented an independent component of the immune response.

### A prolonged hyperinflammatory phase following amputation delays anterior regeneration

In the preceding experiments, we demonstrated that an unresolved immune response caused tissue lysis and that *cathepsin^+^* phagocytes mitigated this cytotoxicity. Given that the endogenous injury-induced inflammatory state normally resolves by 48 hours post-amputation (hpa), prior to the onset of regeneration (Fig. 1A), we hypothesized that persistent inflammation would functionally interfere with the initiation of tissue reconstruction.

To test whether resolving inflammation is a prerequisite for regeneration, we designed an assay to sustain immune activation during the early injury response. We pre-treated intact planarians with a high concentration of heat-killed *E. coli* for 13 hours (Fig. 7A; Methods). We then amputated the animals and immediately returned the tail fragments to the same bacterial solution for an additional 5 hours (Fig. 7A). Subsequently, we transferred the regenerates to a lower bacterial concentration to maintain low-grade immune activation while preserving viability (Fig. 7A; Methods).

**Figure 7.**
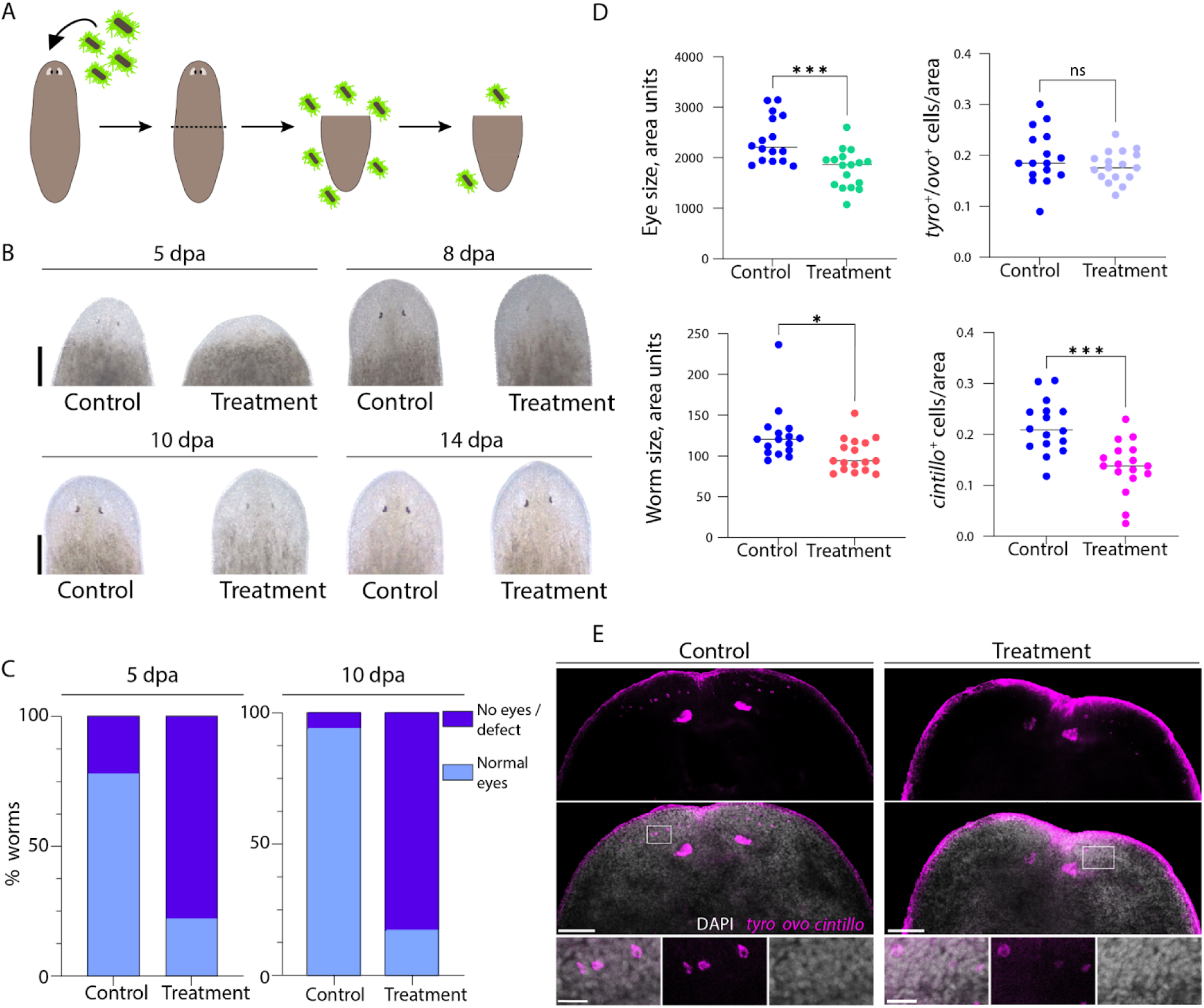
Prolonged and hyperactivated immune response following amputation delays head regeneration. (A) Experimental design. Worms were exposed to a high concentration of heat-killed *E. coli* (∼10^9^ CFU/mL) for 13 hours, amputated, and tail fragments were maintained in the high-concentration solution for 5 additional hours. Subsequently, tail fragments were transferred to a lower concentration (∼6x10^7^ CFU/mL), and regeneration was monitored. (B-C) Regeneration timeline under immune stimulation. (B) Eye development was assessed in control and treated animals from 5 to 14 days post-amputation (dpa) to characterize the observed regeneration delay. At 5 dpa, control animals exhibited pronounced blastemas and pigment development (visible eyespots: n = 14/18), while treated animals showed underdeveloped blastemas, with few animals showing evidence of eyespots (visible eyespots: n = 4/18). By 8 dpa, treated animals developed blastemas but showed minimal eye pigmentation. Control animals were fully regenerated by 10 dpa (n = 17/18), whereas only a minority of the treated animals had completely pigmented eyespots (n = 3/18). By day 14 all animals from both groups completed regeneration. Scale bar: 0.25 mm. (C) Phenotype quantification. Treated animals produced a significantly higher percentage of weak or no-eye phenotypes compared to controls at 5 dpa (22% vs. 78% normal eyes, respectively; p=0.0022) and 10 dpa (17% vs. 94% normal eyes; p= 4. 06 * 10^-6^). P-values were calculated using Fisher’s exact test. (D) Cellular quantification at 5 dpa. Treated worms produced significantly smaller absolute eye areas (top left) and smaller total body sizes (bottom left), reflecting delayed blastema growth. Normalized eye area (*tyro*^+^/*ovo*^+^ cells/area) showed no significant difference between conditions (top right). Treated worms also generated significantly fewer *cintillo*⁺ anterior neurons (bottom right). Horizontal bars indicate the median. (E) Representative FISH images showing differences in eye and anterior neuron development in control vs. treated worms at 5 dpa. Treated worms display smaller eyes, and fewer anterior neurons. Eyes were labeled with *tyrosinase* and *ovo* probes (magenta) and anterior neurons with *cintillo* probes (magenta). Scale: 100 µm. Bottom panels show a magnification of anterior neurons, indicated by white squares. Scale: 25 µm. P-values derived from Fisher’s exact test for categorical data (C) and unpaired two-tailed Student’s t-test for continuous data (D). * p <0.05, *** p <0.001, **** p <0.0001; ns, not significant.

By 5 days post-amputation (dpa), most untreated control fragments developed visible eyespots and distinct blastemas (n=14/18; Fig. 7B-C). By contrast, the majority of treated fragments exhibited weak or absent eye pigmentation and smaller blastemas (n=4/18 developed visible eyespots; Fig. 7B-C). At 10 dpa, control animals displayed fully regenerated heads and eyes (n=17/18), while treated animals continued to lag, displaying reduced or absent eye pigmentation (n=3/18 developed normal eyespots; Fig. 7B-C). By 14 dpa, however, the treated animals successfully formed normal eyes and heads (Fig. 7B). This indicated that sustained immune activation following injury delayed, but did not permanently arrest or dysregulate the regenerative process.

To quantify this developmental delay, we measured eye size and anterior sensory neuron regeneration at 5 dpa using fluorescence in situ hybridization (FISH) (Methods). We labeled eye progenitors and optic-cup cells with *ovo* and *tyrosinase* (*tyro*), respectively ^77,78^, and marked anterior sensory neurons with *cintillo* ^79^ (Fig. 7D-E; Methods). We quantified absolute eye size by measuring the combined area of *ovo*⁺ and *tyro*⁺ cells from both eyes (Methods). Treated planarians possessed significantly smaller absolute eye areas compared to controls (Fig. 7D-E). We also observed a significant reduction in total animal size in the treated group, reflecting their smaller blastemas at this time point (Fig. 7D). When normalized to total animal size, however, the *ovo*⁺*/tyro*⁺ cell area showed no significant difference between the treated and control groups, indicative of a delayed, rather than abnormal regeneration (Fig. 7D-E). Furthermore, treated animals exhibited significantly reduced numbers of *cintillo*⁺ cells, alongside visibly weaker *cintillo* expression (Fig. 7D-E). Taken together, these results demonstrated that maintaining an active pro-inflammatory state interfered with the timely initiation of regeneration, resulting in a developmental delay.

## Discussion

The response to tissue injury requires coordination between host defense and tissue recovery, yet these two demands are not always compatible. While the injury-induced immune response is necessary for immediate survival, it can also create a barrier to tissue replacement ^24,58^. In mammals, excessive inflammatory signaling is proposed to drive fibrosis and scar formation ^23^. Our results indicated that in the highly regenerative planarian *S. mediterranea*, injury likewise activated an innate immune response, however, the inflammatory phase must be brief, and must resolve before the proliferative regenerative phase begins. This demonstrated that resolution of defense responses is a basic evolutionary requirement for a regeneration-permissive environment, such as planarians.

By uncoupling immune activation from physical tissue injury using heat-killed *E. coli*, we found that the planarian generic wound response was intrinsically linked to an innate immune program. The robust induction of early wound response genes, such as *fos-1* and *egrl-1*, together with planarian-specific injury-induced genes such as *hadrian* and *ston*, by non-infective bacterial particles highlighted the deep overlap between pathways that sense tissue disruption and those aimed at responding microbial threats. In this view, the wound response is part of a broader defense state that first stabilizes the organism, with productive tissue replacement delayed until this initial defense response is resolved.

Sustained exposure to non-viable bacteria induced progressive tissue damage and ultimately organismal lysis. Because the bacteria were inactivated, and full recovery of lesions and head regression occurred quickly following the removal of the inactive bacteria, the observed deterioration was most likely host-driven, reflecting a hyperinflammatory state rather than uncontrolled infection. This phenotype was reminiscent of systemic inflammatory pathologies in mammals, such as septic shock, in which self-amplifying immune responses become cytotoxic to the host ^17^. Our findings suggested that planarians were also vulnerable to such uncontrolled inflammatory damage, underscoring why the early immune phase must be rapidly extinguished to prevent host-driven tissue destruction and permit safe tissue repair.

We found that the planarian immune response was dynamic, and appears to require a biphasic cellular response, characterized by an early, predominantly epidermal mediated signaling, and followed by delayed activation of *cathepsin*^+^ phagocytes. This division of labor mirrors a general principle of vertebrate innate immunity, in which barrier tissues initiate the response and phagocytic cells subsequently help restore tissue homeostasis ^11,80,81^. In vertebrates, successful repair depends on the transition of macrophages from pro-inflammatory to pro-resolving states ^27,82^. Although the precise evolutionary relationship between planarian phagocytes and vertebrate immune lineages remains unresolved, members of the Forkhead box (Fox) family are central to both immune resolution and tissue repair in mammals ^83,84^. For example, *foxO1* promotes macrophage phagocytosis and can shift them towards either an anti- or pro-inflammatory phenotype, depending on the activating signals ^85–87^, and *foxF-*1 contributes to mesenchymal tissue repair and limits inflammation ^88^. In this context, our functional data identified planarian *foxF-1*⁺ phagocytes as critical regulators of immune resolution. Suppression of *foxF-1* expression resulted in decreased survival following sustained immune stimulation, suggesting that these cells actively buffer the cytotoxic microenvironment. More broadly, the involvement of a Fox-family transcription factor in this process raised the possibility that regulatory logic linking inflammation resolution to tissue restoration has deep evolutionary roots.

A particularly important observation is that noninfectious immune activation elevated systemic neoblast proliferation while simultaneously delaying regeneration. This uncouples mitotic activation from regenerative success. Inflammatory signaling is well known to act as a potent mitogen in vertebrates, stimulating emergency hematopoiesis and activating tissue-resident stem and progenitor cells ^65,89^. By experimentally sustaining immune activation, we effectively maintained planarian neoblasts in a pro-inflammatory, mitogenic environment. Stem cells were driven to divide, yet this proliferative response did not promote sufficient tissue recovery to overcome the detrimental effects of prolonged bacterial particle exposure. Thus, increased proliferation alone does not ensure regeneration. Instead, our data suggest the observed increase in proliferation constitutes a part of the host response to pathogens, and that regenerative progression depends on whether the inflammatory microenvironment can transition from a defensive to a permissive state.

The evolutionary loss of widespread regeneration in adult mammals is often attributed to the emergence of the adaptive immune system ^33,80^. Our findings raised the possibility that constraints on regeneration may also arise from the dynamics of innate immunity itself. During an active immune challenge, the local tissue environment appears to prioritize defense over repair. The morphogenetic processes required for regeneration are energetically demanding and may be incompatible with a persistent inflammatory state. Comparative models have proposed that endothermic animals prioritize energetic resources for rapid immune defense, potentially at the expense of tissue restoration ^33,90^. Planarians may preserve their exceptional regenerative capacity, at least in part, because they can mount a strong innate immune response and then rapidly resolve it ^35^.

In summary, our findings show a fundamental wiring of immunity into the wound response, and demonstrate that resolution of innate immunity is not merely associated with regeneration, but is a key prerequisite for it. By identifying *foxF-1*⁺ phagocytes as mediators of this transition, we establish planarians as a tractable *in vivo* model for dissecting a broadly relevant checkpoint that separates successful tissue restoration from chronic inflammation and tissue failure.

## Methods

### Animal husbandry

The asexual clonal CIW4 strain of *Schmidtea mediterranea* was used for all experiments. Animals were starved for 7–14 days prior to experiments. Worms were maintained in plastic containers or Petri dishes containing 1x Montjuïc water (i.e. planarian water) (1.6 mmol/L NaCl, 1.0 mmol/L CaCl₂, 1.0 mmol/L MgSO₄, 0.1 mmol/L MgCl₂, 0.1 mmol/L KCl, and 1.2 mmol/L NaHCO₃ prepared in Milli-Q water) and housed at 20°C under dark conditions. Cultures were fed blended calf liver once weekly and cleaned twice per week.

### Live and heat-killed bacteria culturing for immune activation assays

Live and heat-killed preparations of the following bacterial strains were used: *E. coli* TOP10, *B. subtilis* PY79, and *P. aeruginosa* PAO1. Bacteria were cultured in Luria-Bertani (LB) medium (1% tryptone (Neogen, CAT. #NCM0211A), 0.5% yeast extract (Gibco, CAT. #212750), and 1% NaCl (Biolab, CAT. #001903059100)) at 37 °C until reaching an OD₆₀₀ of approximately 1. Cultures were then pelleted and cells were washed three times with planarian water.

For heat inactivation, bacterial suspensions were incubated in a heat block at 95 °C for 15 min (*E. coli*) or at 85 °C for 10 min (*B. subtilis* and *P. aeruginosa*). To confirm successful inactivation, undiluted samples were plated on LB agar plates; no colony growth was observed following heat treatment. Prior to heat inactivation, dilutions of live bacteria were plated on LB plates and colonies were counted to determine bacterial concentration.

### Preparation of FITC-labeled heat-killed *E. coli*

Heat-killed *E. coli* were pelleted and resuspended in a 1 mg/mL solution of fluorescein isothiocyanate (FITC) (Sigma-Aldrich, CAT. #F6377) in 0.05 M carbonate buffer (pH 9.3) to a concentration of 10^8^ - 10^9^ CFU/mL. The inactivated bacteria were incubated with the FITC solution for 1 hour at room temperature in the dark. The FITC-labeled cultures were subsequently washed three times by pelleting and resuspending the inactivated bacteria in planarian water.

### RNA isolation for sequencing and RT-qPCR

Samples were collected into 700 μL TRI Reagent (Sigma; CAT. #9424) and then homogenized using 0.5 mm zirconium beads in a bead-beating homogenizer (Allsheng; Bioprep-24) for two cycles of 45 s at 3,500 rpm followed by a 5-min incubation at room temperature. Then, 150 μL of chloroform was added to each tube. Tubes were shaken for 15 s and then incubated at room temperature for 3 min. Then, samples were centrifuged at 12,000 g for 30 min at 4 °C, and the aqueous upper phase was transferred to a new tube. Isopropanol (500 μL) was added, and tubes were inverted five times followed by a 10-min incubation at room temperature. Samples were centrifuged at 4°C (12,000 *g*) for 45 min. The supernatant was removed and the pellet was washed twice with 75% EtOH, centrifuged at 7,500 *g* for 5 min at 4°C, and air-dried for 10 min. RNA was resuspended in 30 μL of nuclease-free water. RNA concentration was measured by Qubit Fluorometer (Invitrogen; CAT. #Q33226) according to the manufacturer’s protocol.

### Double-stranded RNA synthesis for RNAi experiments

Double-stranded RNA (dsRNA) was synthesized as previously described ^91^. Briefly, *in vitro* transcription (IVT) templates were prepared by PCR amplification of cloned target sequences using primers with 5ʹ flanking T7 promoter sequences. dsRNA was synthesized using the TranscriptAid T7 High Yield Transcription Kit (CAT. #K0441; Thermo Fisher Scientific). Reactions were incubated overnight at 37°C and then supplemented with RNase-free DNase I for 30 min. RNA was purified by ethanol precipitation and finally resuspended in 30 μL of ddH₂O. RNA was analyzed on a 1% agarose gel and quantified by Qubit 4 (Thermo Fisher Scientific) to confirm concentrations higher than 5 μg/μL. Animals were starved for at least 7 days prior to RNAi experiments. Asexual *Schmidtea mediterranea* were used for all experiments. Animals were fed with dsRNA mixed with beef liver twice a week as previously described ^92^. For *foxF-1* RNAi experiments, animals were fed dsRNA twice, and processed for RNA isolation or fixation 9-10 days post-last feeding. The non-planarian gene *unc22* was used as control dsRNA for all RNAi experiments.

### cDNA synthesis for RT-qPCR

RNA isolated from samples was converted to cDNA using RevertAid H Minus First Strand cDNA Synthesis Kit (Thermo Fisher Scientific; CAT: #K1631). Oligo(dT) primers were used to obtain cDNA from polyadenylated RNA. The resulting cDNA was used for qPCR analysis.

### Gene expression quantification by RT-qPCR

2x qPCRBIO Fast qPCR SyGreen Blue Mix (PCR Biosystems; CAT. #PB20.15-20) was used for all RT-qPCR experiments. The expression of the target genes was measured using the QuantStudio 3 Real-Time PCR system (Applied Biosystems) with the following program: [95°C for 20 seconds, 40 cycles (95°C for 1 second, 60°C for 20 seconds)]. At least two biological replicates were used per condition, with two technical replicates per sample. The relative gene expression fold-change was calculated using the ΔΔCt method, with *gapdh* used as an endogenous control. Forward and reverse primers were designed for a 90-150 bp section for each target gene. Primer efficiency was tested prior to experiments using a standard curve generated from four serial dilutions of cDNA.

### Preparation of RNA sequencing libraries following exposure to heat-killed *E. coli*

Worms measuring 5-6 mm in length were incubated with heat-killed *E. coli* (∼10^9^ CFU/mL) for 20 and 40 hours. Control groups not exposed to bacteria were isolated at each time point. For each RNA-seq library, 1 μg of purified total RNA was used. Illumina-sequencing compatible cDNA libraries were prepared using NEBNext Ultra II Directional RNA Library Prep Kit for Illumina (New England Biolabs; CAT. #E7760L), according to the manufacturer’s protocol. Briefly, mRNA was enriched by poly(A) selection using oligo(dT) magnetic beads. Then, RNA was fragmented according to the protocol, and complementary DNA (cDNA) was produced by reverse transcription. Following second-strand synthesis, the resultant double-strand DNA (dsDNA) was end-repaired, and then adaptor ligation was performed according to protocol. The libraries were then amplified by PCR using barcoded primers according to the kit instructions.

### Analysis of RNA sequencing gene expression changes

Sequenced reads were aligned to the *S. mediterranea* reference transcriptome (assembly dd_Smed_v6 ^93^) using Bowtie2 (version 2.4.1) ^94^, with trimming of 10 nt from the 3’ ends of the reads. The resulting SAM alignment files were subsequently converted to binary format and sorted using Samtools (version 1.19). Following alignment, read quantification was performed to generate a raw gene expression matrix using featureCounts (version 2.0.0) with parameters [-M -s 0]. Differential expression analysis was then conducted with the DESeq2 package ^95^ using pairwise statistical contrasts to identify differentially expressed genes between each treatment and its respective control.

### Analysis of planarian injury-time course gene expression data

Pre-computed differential expression files of injury time-course ^41^ between each time point and its control were downloaded from PlanaTools ^96^ along with a gene expression matrix. Individual differential expression tables were integrated into a single differential expression time-course dataset. Significantly upregulated genes (Fold change > 2, FDR < 0.05) were selected for clustering analysis. Gene expression data across all time points, excluding the 48 hours post-amputation (hpa) samples, were normalized using row-wise z-scores. k-means clustering was performed after examining the optimal cluster numbers using the elbow method. The k-means approach was applied using 25 random initial configurations (nstart = 25) and a maximum of 100 iterations (iter.max = 100) to ensure convergence and determine the final cluster assignments.

### Gene cloning and transformation

Genes were amplified from planarian cDNA using gene-specific primers and subsequently cloned into the pGEM-T vector according to the manufacturer’s instructions (Promega; CAT. #A1360) (Table S6). Primers were designed to include standard pGEM-T compatible overhang sequences (Forward: 5ʹ-AAGCTGGAGCTCCACCGCGG-3ʹ; Reverse: 5ʹ-GGGCGAATTGGGTACCGGG-3ʹ). Plasmids containing cloned *cintillo* and *CSTL2* (*dd_175*) sequences were kindly provided by the Reddien Lab.

Recombinant vectors were transformed into *E. coli* TOP10 cells (Thermo Fisher Scientific) using the heat-shock method. Briefly, 100 μL of bacteria was mixed with 5 μL of each of the cloned vectors, incubated on ice for 30 min, and then placed at 42°C for 45 s. Then, the transformed bacteria were supplemented with 350 μL LB medium, and following 1 h of recovery at 37°C, the bacteria were plated on agar plates containing ampicillin (1:1000), isopropyl β-D-1-thiogalactoside (IPTG) (1:1000), and 5-bromo-4-chloro-3-indolyl-β-D-galactopyranoside (X-gal) (1:500). Colonies were grown overnight at 37°C, and colonies were screened by colony PCR using CPC21 and CPC22 primers with the following PCR program: (i) 5 min at 95°C; (ii) 34 cycles of 45 s at 95°C, 60 s at 55°C, and 2:30 min at 72°C; (iii) 10 min at 72°C; (iv) hold at 10°C. Reactions were analyzed by gel electrophoresis, and correctly sized gene products were grown overnight in LB medium, supplemented with ampicillin (1:1000) at 37°C. Plasmids were purified from overnight cultures with the NucleoSpin Plasmid Miniprep Kit (Macherey-Nagel; CAT. #740588). Cloned gene sequences were sequenced by Sanger sequencing.

### Fixation for whole-mount immunofluorescence and FISH

Fixation was performed as previously described ^97^. Animals were placed in 5% N-Acetyl-L-cysteine (NAC) (Mercury; CAT. #1124220100) in PBS for 5 min, then incubated in 4% formaldehyde (FA) in 0.1% PBSTx (phosphate-buffered saline, 0.1% Triton X-100) for 20 min. Animals were then washed in PBSTx, 50:50 PBSTx:methanol and stored in methanol at -20°C.

### Whole-mount fluorescence in situ hybridization (FISH)

Fluorescence in situ hybridization (FISH) was performed as previously described ^97^ with minor modifications. Briefly, fixed animals were bleached and treated with proteinase K (2 μg/mL; Thermo Fisher Scientific; CAT. #25530049) in PBSTx. Following overnight hybridizations, samples were washed twice in pre-hyb solution, 1:1 pre-hyb:2X SSC, 2X SSC, 0.2X SSC, PBSTx. Subsequently, blocking was performed in 0.5% Roche Western Blocking Reagent and 5% inactivated horse serum (HIHS) in PBSTx. Animals were incubated overnight at 4°C with anti-DIG-POD antibody (Roche; CAT. #11207733910) diluted 1:1,500 in blocking solution. Post-antibody washes and tyramide development were performed as previously described ^97^. Specimens were counterstained with DAPI overnight at 4°C (1 μg/mL in PBSTx), and mounted on slides using Vectashield antifade mounting media (Vector Laboratories; CAT. #H-1000-10)

### Immunofluorescence for *foxF-1*-controlled muscle labeling

Following 4% FA fixation, worms were bleached and treated with proteinase K (2 μg/mL) in PBSTx, followed by 1-2 h blocking as described previously. Worms were then incubated with anti-muscle 6G10 (Mouse) antibody solution (https://dshb.biology.uiowa.edu/, RRID: https://www.antibodyregistry.org/AB_2619613) overnight at 4°C.

### H3P labeling

For H3P labeling, animals were fixed in 2% HCl and placed on ice for 30 seconds, then transferred to Carnoy’s fixative (60% ethanol, 30% chloroform, 10% glacial acetic acid) for 5 minutes at room temperature, then for another 2 hours on ice ^98^. H3P labeling was performed based on published protocols ^99^ ^54^. Briefly, following fixation, bleaching was performed in 6% hydrogen peroxide (H_2_O_2_) in methanol overnight on a light table at room temperature. Subsequently, blocking was performed with 10% heat inactivated horse serum in PBSTx. Samples were incubated with anti-phospho-histone H3 antibody (anti-H3P)(1:100; Sigma-Aldrich; CAT. #04-817) overnight at 4°C. Anti-H3P antibody was then washed seven times with PBSTx for 20 min, and subsequently incubated with goat anti-rabbit-HRP secondary antibody (1:100; Abcam; CAT. #ab6721) overnight at 4°C. The secondary antibody was washed seven times and samples were developed with rhodamine tyramide diluted 1:1,000 in PBSTi. Samples were then mounted on slides using Vectashield antifade mounting media (Vector Laboratories; CAT. #H-1000-10).

### Imaging and cell counting

Fluorescence and confocal images of samples labeled with FISH, immunofluorescence and fluorescent bacteria were acquired using a confocal microscope (Zeiss LSM 800). Live images were taken using a stereomicroscope (Leica S9i). Labeled cells were counted manually using the cell counter module in the ImageJ software ^100^. H3P⁺ cells were counted throughout the entire animal; *CSTL2*⁺ cells were counted in a rectangle between the anterior end of the pharynx and the posterior end of the brain. Normalization of H3P⁺/ *CSTL2*⁺ cells was performed by dividing the number of labeled cells by the area that was counted.

### Statistical analysis

Statistical analysis was performed using GraphPad Prism v10.6.1 or R 4.3. Comparison between two groups was performed by unpaired Student’s t-test, unless stated otherwise. Comparison between more than two groups was performed by one-way ANOVA followed by Holm-Šídák multiple comparisons test where applicable, unless stated otherwise.

### Eye size measurement and *cintillo*+ neuron quantification

Eye size was assessed by measuring the area of *ovo*- and *tyrosinase*-labeled cells in each eye using the area-measurement module of ImageJ^100^. Combined eye area (left and right eyes) was then plotted for quantification. Normalization of eye area was performed by dividing eye area by area of the entire regenerate at 5 dpa. Worm area was measured by the circumference of the largest z-slice. Anterior neuron quantification was done by manually counting *cintillo*^+^ cells using the ImageJ cell counter module.

### Preparation of scRNAseq libraries following exposure to heat-killed *E. coli*

Worms sized 5-6 mm were incubated with heat-killed *E. coli* (∼10^9^ CFU/mL) for 40 hours. Bacteria were then washed and worms were macerated into small fragments using a scalpel. Tissue fragments were collected into calcium-free, magnesium-free medium plus 0.1% BSA (CMFB) and dissociated by pipetting for 5 minutes and dissociated cells were strained through a 40 μm filter. Cells were centrifuged at 1250 rpm for 5 min at 4°C and resuspended in CMFB containing Hoechst 33342 (20 μL/mL) for 30 min at room temperature, in the dark. Before loading into BD FACSymphony S6, cells were labeled with propidium iodide (5 μg/mL) for cell viability detection. FACS gating was performed as previously described for planarian cell populations ^101^. Viable cells were isolated and immediately processed for single-cell library preparation using 10X Genomics GEM-X OCM 3’ Chip Kit (CAT. #1000747) having two replicates per condition (e.g., exposed to heat-killed bacteria and unexposed controls). scRNA-seq libraries were sequenced on Illumina NextSeq 2000 at The Rosalie and Harold Rae Brown Cancer Research Core Facility at the Faculty of Life Sciences in Tel Aviv University according to the manufacturer’s protocol.

### Classification of immune related genes

Differentially expressed genes associated with planarian injury and host responses were classified as immune-related based on their top BLAST annotations (Tables S1–S2). Only genes with assigned annotations were included in this classification. These genes were further categorized as either immune activators or immune suppressors based on published functional evidence, considering their reported roles in relevant biological contexts such as injury and immune induction or infection.

### Analysis of host response single-cell RNAseq

scRNA-seq count matrices from four samples (two controls, two heat-killed bacteria-treated) were imported and processed using Seurat ^71^. Cells expressing 200 - 7,000 detected genes and at least 500 unique molecular identifiers (UMIs) were retained for further purposes. Samples were merged and the integrated dataset was log-normalized. The top 2,000 highly variable features were selected, scaled, and ribosomal transcript percentage was regressed out. Principal Component Analysis (PCA) was performed, and first 35 principal components were used for performing clustering (resolution 0.5), and generating UMAP and t-SNE embeddings for visualization. Cluster-specific marker genes were identified using the Seurat FindAllMarkers function, applying a Wilcoxon rank-sum test with parameters [only.pos = TRUE, min.pct = 0.25, logfc.threshold = 0.25]. To evaluate the transcriptional response to the bacterial treatment, differential gene expression between treated and control cells within each cluster was assessed using the Seurat FindMarkers function with parameters [min.pct = 0.1, logfc.threshold = 0.1]. Finally, Fisher’s exact test was utilized to identify statistically significant shifts in the proportional representation of cells within each cluster across the two treatment conditions with p-value adjusted by the Benjamini-Hochberg method. Quantification of immune-related gene set activity enrichment scores at single-cell resolution were calculated using the UCell package ^70^. A manually curated list of planarian immune-related genes (Table S1) was utilized to score individual cells within the integrated Seurat object using the AddModuleScore_UCell function. These UCell enrichment scores were subsequently compared between heat-killed bacteria-treated and control samples across the identified clusters. Statistical significance was determined using a Wilcoxon rank-sum test, and resulting p-values were adjusted for multiple testing using the Benjamini-Hochberg correction.

## Data availability

Sequencing data produced here, including bulk- and single-cell RNA-seq, is to be released via the Sequence Read Archive (SRA) ^102^ with BioProject accession PRJNA1466577.

## Acknowledgments

We thank Prof. Benyamin Rosental for his valuable discussions and feedback during this work. We are grateful to Prof. Avigdor Eldar, Dr. Shira Omer Bendori, Dr. Nitzan Aframian for their assistance with the experiments involving *B. subtilis* and *P. aeruginosa*. We thank Prof. Eliora Ron for her valuable discussions. We acknowledge Dr. Hila Kobo and Dr. Tamar Katzir from the Tel Aviv University Genomics Unit, Faculty of Life Sciences Cancer Research Core Facility (CRCF), for their assistance with RNA-seq library preparation and Illumina sequencing at the Faculty of Life Sciences core facilities. We thank Dr. Rami Khosravi for assistance with scRNA-seq library preparation at the Faculty of Medicine Research Infrastructure Core Facilities (RICF), Tel Aviv University. We thank Prakash Varkey Cherian, Tamar Zelig, and members of the Wurtzel laboratory for their assistance and feedback on the manuscript. O.W. is supported by the Israel Science Foundation (no. 2741/25) and the European Research Council (no. 853640). O.W. is a Zuckerman Faculty Scholar.

## Supplementary Figures

**Figure S1.**
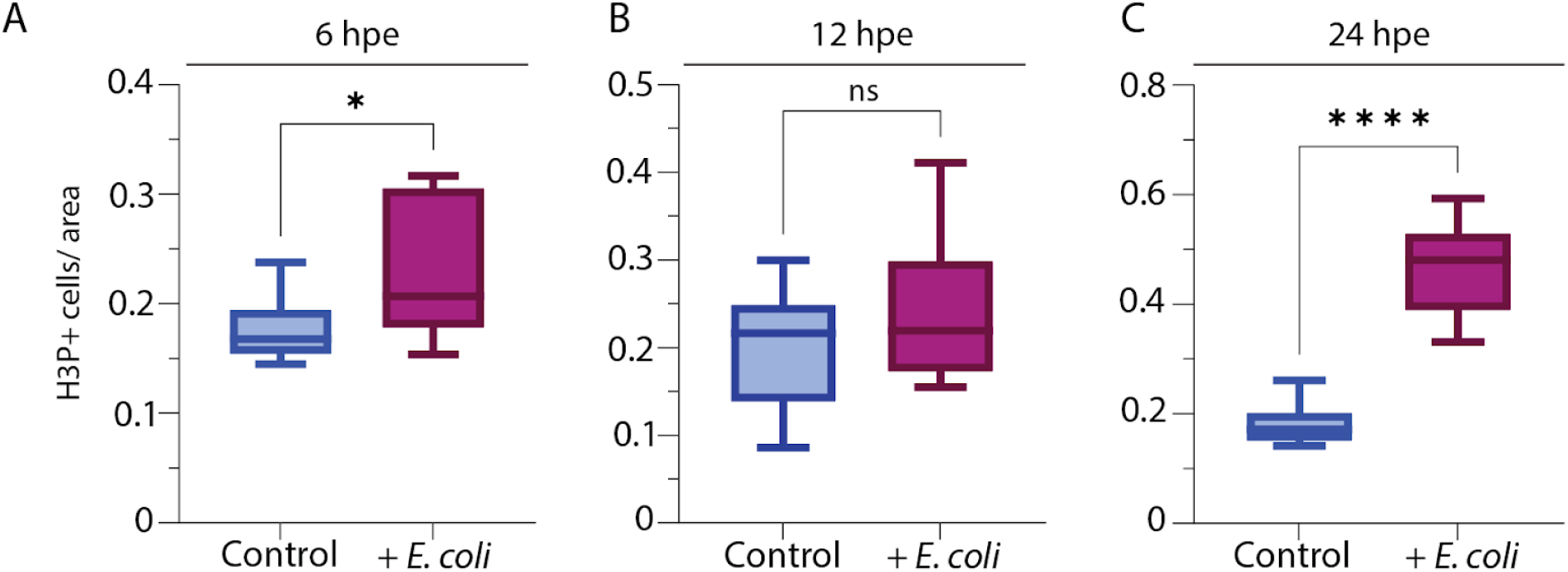
Exposure to heat-killed *E. coli* induces two mitotic peaks in planarians. (A-C) Plots showing normalized H3P⁺ counts (Methods), at 6, 12 and 24 hours post-exposure (hpe). For 6 hpe control: n=11; for 6 hpe treated: n=11; for 12 hpe control: n=12; for 12 hpe treated: n=10; for 24 hpe control: n=13; for 24 hpe treated: n=13. P-values were calculated using unpaired two-tailed Student’s t-test (*** p<0.0001, * p<0.05), horizontal lines indicate median, error bars indicate minimal and maximal values.

**Figure S2.**
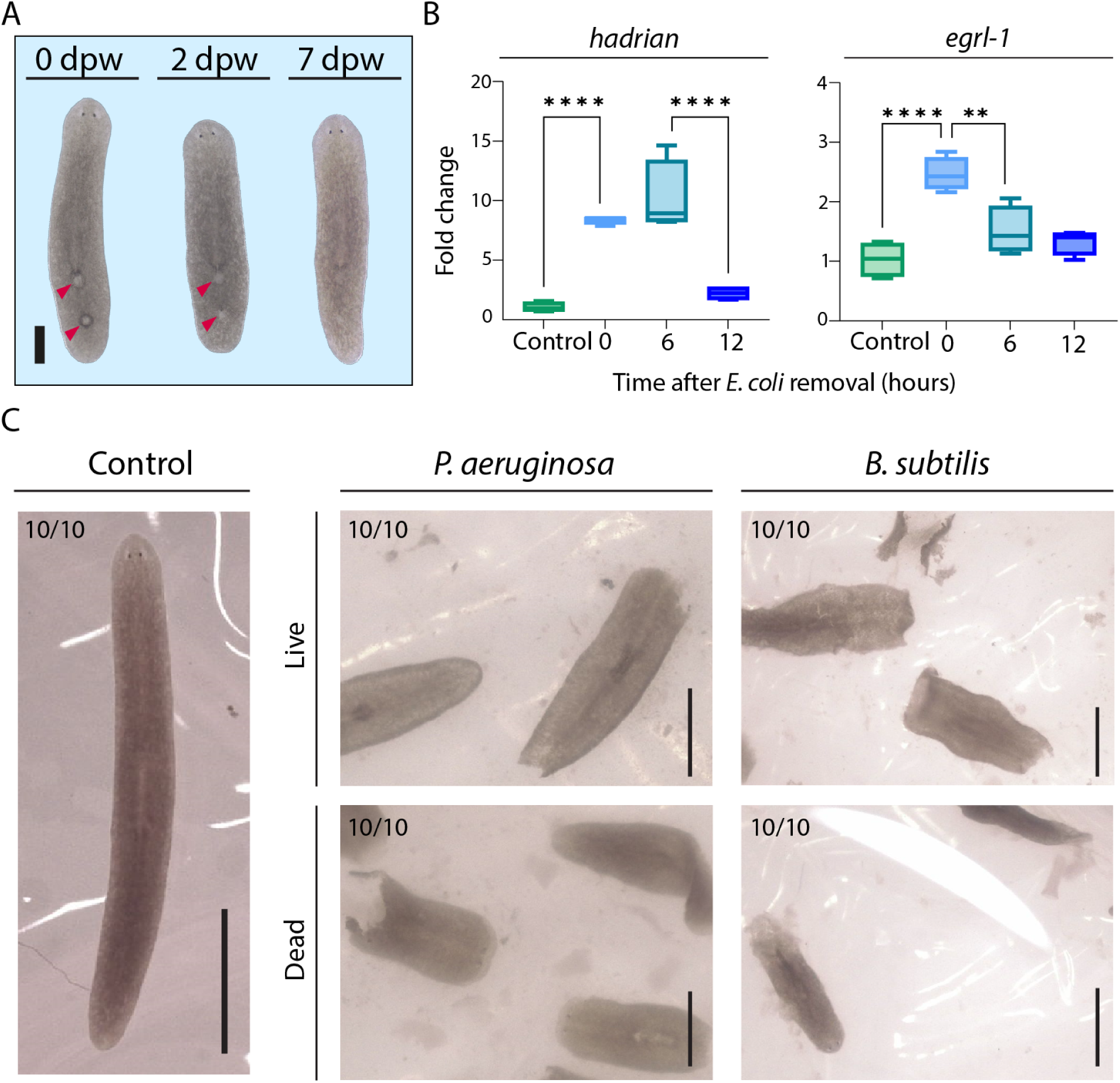
Heat-killed bacteria induce a reversible injury-like response associated with tissue damage and lysis. (A) Dorsal lesions heal by regeneration following removal of heat-killed bacteria. From left to right: two dorsal lesions (indicated by red arrows) at 0 days post-bacterial wash (dpw); the lesions after 2 dpw; lesions healed almost completely by 7 dpw. (B) RT-qPCR analysis of *hadrian* and *egrl-1* gene expression changes following removal of heat-killed *E. coli* from the environment; Y-axis indicates fold change; X-axis indicates time after removal of *E. coli* in hours; *hadrian* expression remained high at 6 hours post-bacterial wash (hpw), and decreased to near-control level by 12 hpw (p=ns); *egrl-1* expression decreased rapidly following removal of bacteria, decreasing to near-control levels by 6 hpw (unchanged by 12 hpw, p=ns, not significant); only significant p-values are indicated. P-values were calculated by ordinary one-way ANOVA, followed by Holm-Šídák’s multiple comparisons test (**** p <0.0001, ** p<0.01). Horizontal line indicates the median, error bars indicate minimum and maximum values. (C) Exposure to *Bacillus subtilis* and *Pseudomonas aeruginosa* caused an identical phenotype as exposure to *E. coli*. From left to right: unexposed animals (control) remained viable throughout the experiment; all worms exposed to live (top) and heat-killed (bottom) *B. subtilis* and *P. aeruginosa* deteriorated. Lysis occurred in 100% of cases, following exposure to either live or heat-killed *B. subtilis* and *P. aeruginosa*; time until lysis varied between groups, on average worms died 1-2 days post exposure to either live or dead *B. subtilis* and *P. aeruginosa*; bacterial concentration used: ∼2-5x 10^8^ CFU/mL.

**Figure S3.**
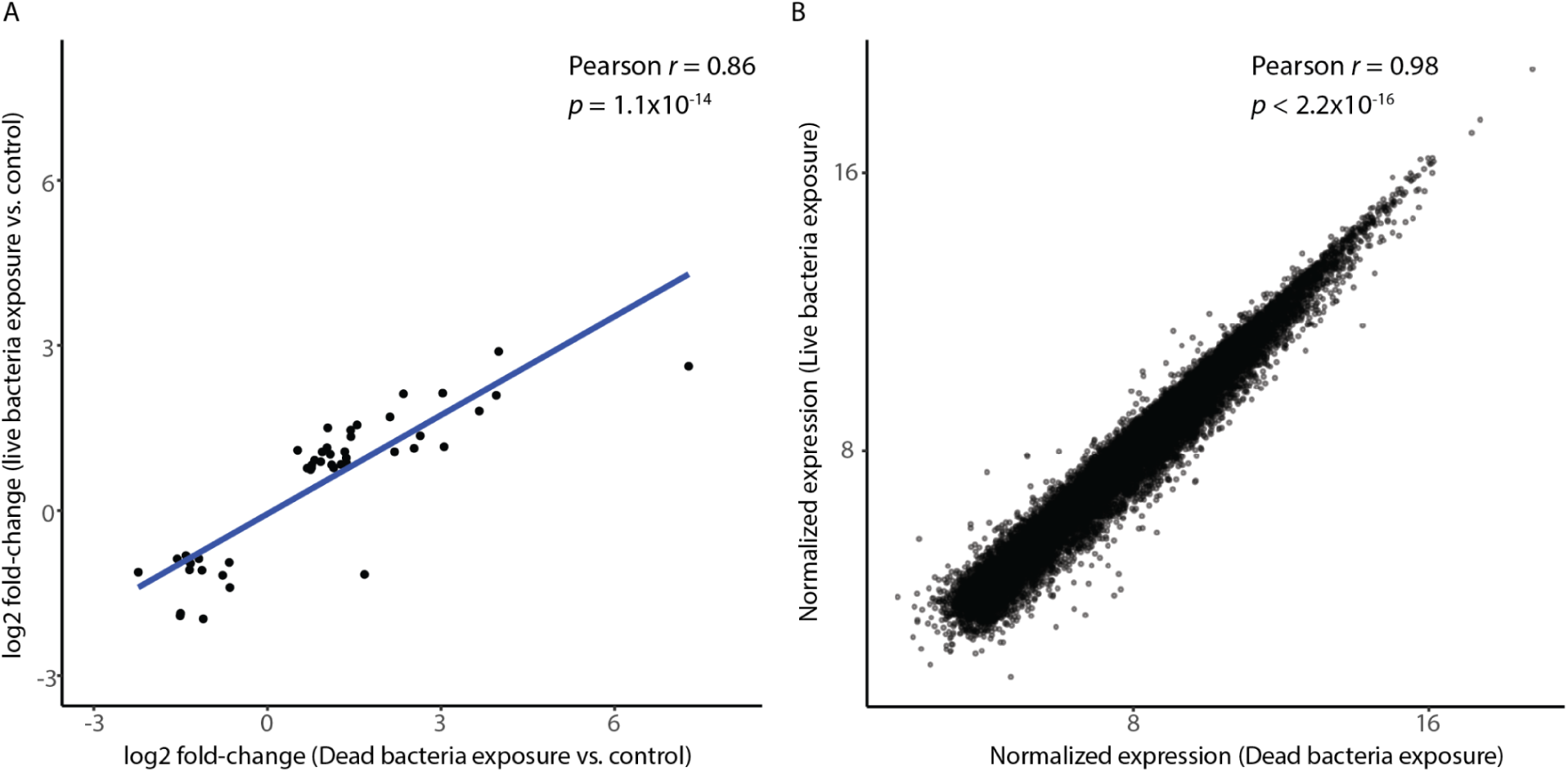
Exposure to live or dead bacteria elicits similar transcriptional responses. (A) Correlation of differentially expressed genes between the two libraries. Values represent the log fold change in gene expression for the experimental samples compared to their respective controls (x and y axes, exposure to dead and live bacteria, respectively). Each dot represents a differentially expressed gene (adjusted *p* < 0.1). (B) Scatter plot comparing average gene expression in planarians exposed to dead versus live bacteria. Each dot represents an individual gene plotted by its average Variance Stabilizing Transformation (VST)-normalized expression (calculated using DESeq2 ^95^. The x-axis denotes expression in planarians exposed to dead bacteria, and the y-axis denotes expression in planarians exposed to live bacteria.

**Figure S4.**
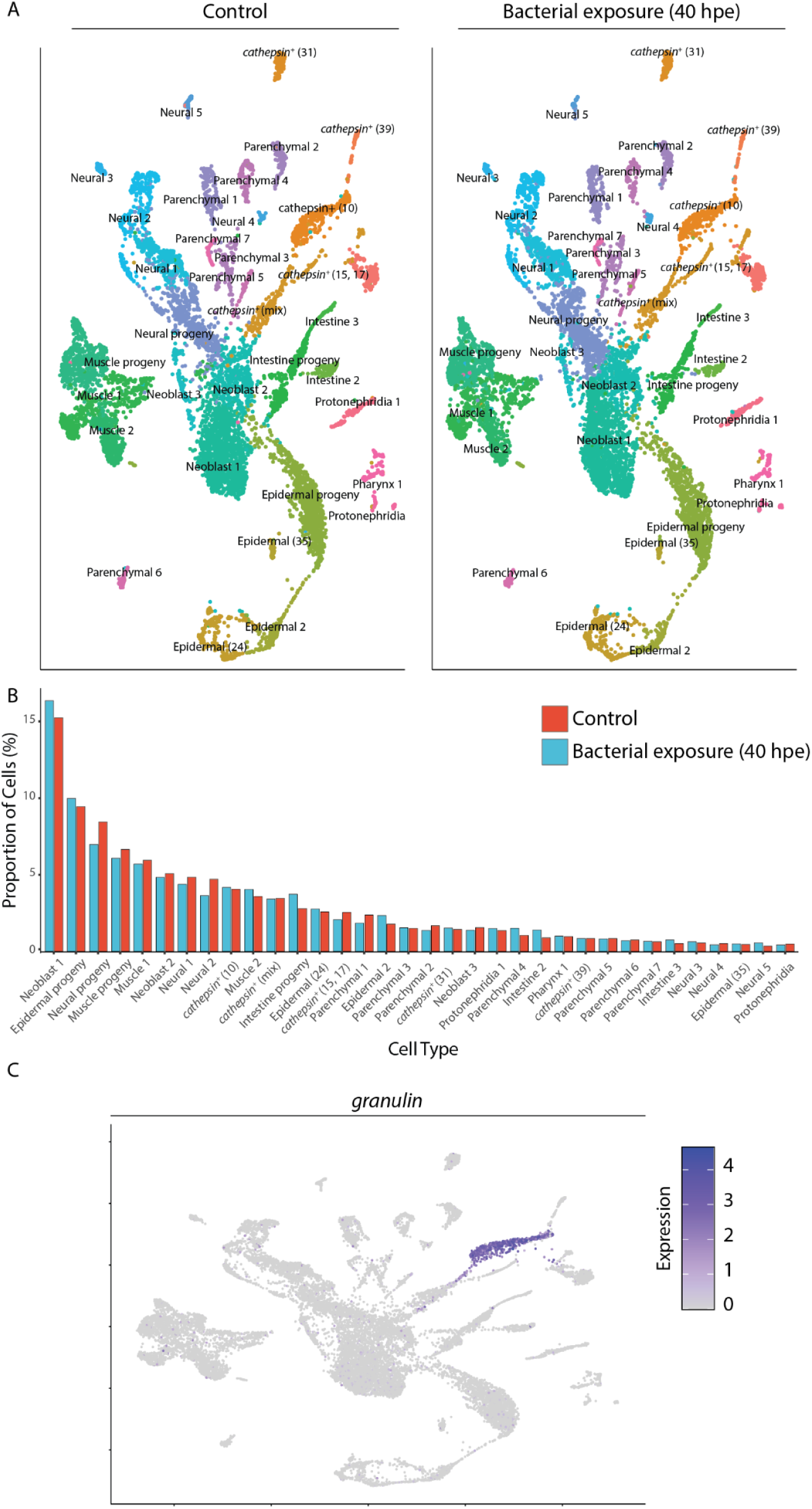
scRNA-seq analysis of the planarian immune response. (A) UMAP plot visualizing single-cell transcriptomic data from planarians under control (unexposed) and 40 hours post-exposure (hpe) to heat-killed *E. coli.* Distinct cell populations are color-coded and annotated by cell type and lineage (e.g., Neoblast, Neural, Epidermal, Parenchymal, and *cathepsin*^+^ clusters), according to the planarian single-cell atlas ^57^. (B) Bar graph quantifying the relative proportion of each annotated cell type as a percentage of the total cell population between the control group (red) and the heat-killed bacterial exposure group at 40 hpe (blue). The similarity between the two groups indicated that the transcriptional responses observed at this timepoint were more compatible with changes to gene expression rather than the enrichment or depletion of specific cell populations. (C) The expression of *granulin* was overlaid on a UMAP plot. The color gradient indicates relative expression levels (gray to purple, low to high expression, respectively).

**Figure S5.**
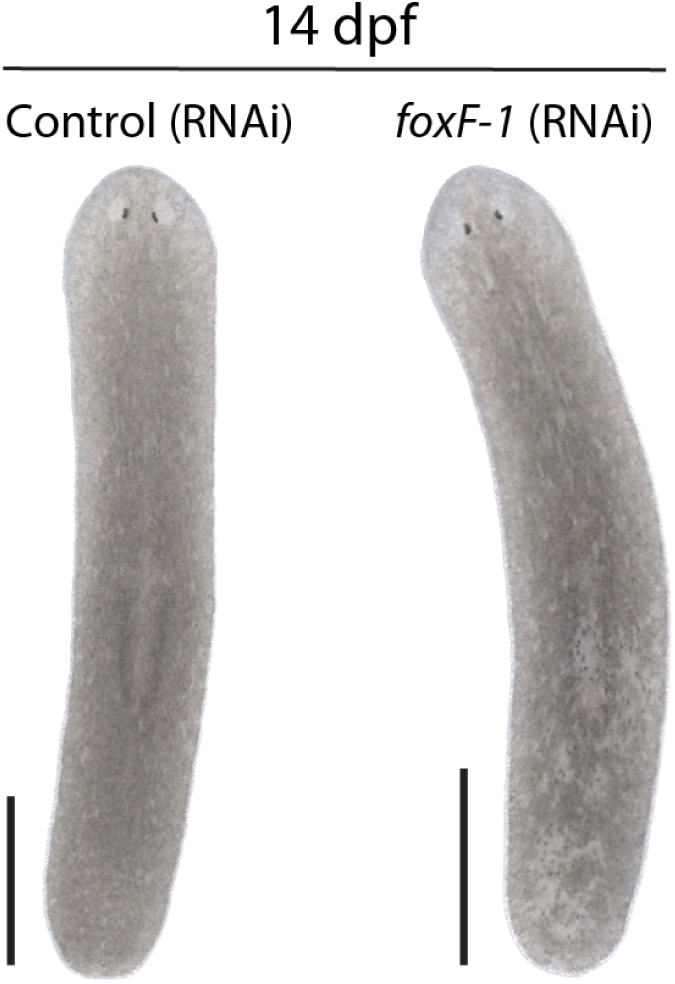
*foxF-1* (RNAi) animals are viable two weeks after *foxF-1* suppression. Representative images of control (left; n=12) and *foxF-1* (RNAi) (right; n=12) animals at 14 days post-last feeding (dpf) after two dsRNA feedings (Methods). Scale: 1 mm.

## Supplementary Tables

**Supplementary Table S1.** Injury-induced gene expression dynamics post-amputation.

**Supplementary Table S2.** Differential gene expression analysis of the planarian host response to heat-killed *E. coli* at 20 and 40 hours post-exposure.

**Supplementary Table S3.** Differential gene expression analysis of the planarian host response at 40 hours following exposure to live *E. coli*.

**Supplementary Table S4**. Differentially expressed marker genes across cell clusters.

**Supplementary Table S5.** Differential expression of cluster marker genes following exposure to heat-killed *E. coli* compared to controls.

**Supplementary Table S6.** Primer sequences used in this study.

